# The virtual aging brain: a model-driven explanation for cognitive decline in older subjects

**DOI:** 10.1101/2022.02.17.480902

**Authors:** Mario Lavanga, Johanna Stumme, Bahar Hazal Yalcinkaya, Jan Fousek, Christiane Jockwitz, Hiba Sheheitli, Nora Bittner, Meysam Hashemi, Spase Petkoski, Svenja Caspers, Viktor Jirsa

**Author notes:** These authors contributed equally (Mario Lavanga, Johanna Stumme, Svenja Caspers, Viktor Jirsa). Corresponding authors: ML, VJ.

## Abstract

Healthy aging is accompanied by heterogeneous decline of cognitive abilities among individuals, especially during senescence. The mechanisms of this variability are not understood, but have been associated with the reorganization of white matter fiber tracts and the functional co-activations of brain regions. Here, we built a causal inference framework to provide mechanistic insight into the link between structural connectivity and brain function, informed by brain imaging data and network modeling. By applying various degrees of interhemispheric degradation of structural connectivity, we were not only able to reproduce the age-related decline in interhemispheric functional communication and the associated dynamical flexibility, but we obtained an increase of global modulation of structural connectivity over the brain function during senescence. Notably, the increase in modulation between structural connectivity and brian function was higher in magnitude and steeper in its increase in older adults with poor cognitive performance. We independently validated the causal hypothesis of our framework via a Bayesian approach based on deep-learning. The current results might be the first mechanistic demonstration of dedifferentiation and scaffolding during aging leading to cognitive decline demonstrated in a large cohort.

## Introduction

Healthy aging is accompanied by a decline of cognitive abilities with substantial variations among the individual aging trajectories, in particular at later stages in life (Hedden & Gabrieli, 2004; Oschwald et al., 2020). The mechanisms of this variability are not understood, but have been associated with organizational changes in structure and function, notably structural connectivity (SC), i.e. white-matter fiber tracts connecting gray matter brain regions (Piguet et al., 2009), and functional connectivity (FC), i.e. co-activation of brain regions (Bassett & Sporns, 2017;Straathof et al., 2019).

As fiber connections are expected to deteriorate (Antonenko & Flöel, 2014; Damoiseaux, 2017; Zuo et al., 2017), particularly with respect to the number of inter-hemispheric fibers within tracts and fiber density (Puxeddu et al., 2020), the subsequent consequences on the organization of functional networks remain elusive. Research on the subject has been restricted to the sole FC to express brain dynamics and to linear-correlation studies, which found a positive and increasingly stronger SC-FC relation in the older population (Fukushima et al., 2018; Greicius et al., 2009; Honey et al., 2009; Jung et al., 2017; Wendelken et al., 2017; Zimmermann et al., 2016). Moreover, empirical findings show that FC and SC independently change during aging (Fjell et al., 2017) and FC might only partially mimic underlying SC (Suárez et al., 2020; Tsang et al., 2017). A one-to-one mapping will not sufficiently describe the dependency between SC and the empirical FC (Damoiseaux, 2017; Mollink et al., 2019; Roland et al., 2017; Straathof et al., 2019), especially at the individual level (Zimmermann et al., 2019). These findings are consistent with nonlinear mappings between SC and functional metrics including descriptors of functional connectivity dynamics (FCD) (Battaglia et al., 2020; Hansen et al., 2015), which capture functional higher-order features such as non-stationarity.

Since degradation of healthy brain structures occur even in the absence of a neuropathological process, such as cellular (myelin pallor, gliosis) and vascular abnormalities (increased perivascular space, reduced perfusion) (Piguet et al., 2009) and it seems to especially take place at the interhemispheric level (Mollink et al., 2019; Puxeddu et al., 2020; Roland et al., 2017), one may suggest a causal hypothesis of interhemispheric SC deterioration which could lead to an overall functional reorganization and its associated cognitive decline. How this causal tethering might unfold in senescence is yet to be explained.

Biophysically realistic network modeling investigations demonstrated that a high similarity of SC and FC is a trivial consequence of linear network dynamics and characteristic of a brain systemy (Saggio et al., 2016). The dynamic flexibility or brain fluidity is expressed by dynamic changes in FC over time (functional connectivity dynamics (FCD)), which has been reported empirically (Lurie et al., 2020) and captures the temporal variations of resting-state (Battaglia et al., 2020). FCD is an indicator of the brain’s capacity to explore flexibly a large range of brain states, which reduces with cognitive decline in both healthy aging (Battaglia et al., 2020) and Alzheimer’s disease (Córdova-Palomera et al., 2017). Additionally, there is a growing body of literature that observed evidence of a nonlinear mapping between SC and FC (John et al., 2022; Shine et al., 2019) and especially between interhemispheric SC and homotopic FC (Mollink et al., 2019; Roland et al., 2017), between SC and FCD (John et al., 2022; Naik et al., 2017; Shen et al., 2015) as well as respective age-related findings (Esfahlani et al., 2021; Naik et al., 2017; Roland et al., 2017), which might benefit from a more holistic approach based on modeling to explain the link between structure and function. To deepen the understanding of the complex link between structural changes, functional changes and cognitive decline, the current challenge for aging models is then twofold: on the one hand, the comprehension of the cause-effect of SC and its functional phenotype and on the other, the link to cognitive decline as suggested by various aging theories (Goh, 2011; Naik et al., 2017; Reuter-Lorenz & Park, 2014). Employing biophysical models accounting for the nonlinearity and the causality of this relationship as well as the wider nature of FC appears highly promising, especially at the individual level.

To shed light on the complex relation between structural connectivity and brain function and its association with cognitive decline for each individual age trajectory, we developed a causal inference framework. Changes in SC are mechanistically expressed in a computational brain model framework, The Virtual Brain (TVB) (Sanz Leon et al., 2013). Virtual brains are data-driven brain network models (Bassett et al., 2018; Breakspear, 2017; Deco et al., 2011; Ghosh et al., 2008; Sanz Leon et al., 2013), integrating individual brain imaging data on brain structure (anatomy, connectivity) to constrain network parameters. Brain network models can simulate resting-state functional data with a high degree of dynamical flexibility and investigate the role of different anatomical connections for generating functional co-activations, such as the interhemispheric tracts or the links belonging to a specific region (Straathof et al., 2019). By modifying the SCs, such as decreasing magnitude of connection strengths along relevant tracts (Deco et al., 2021; Messé et al., 2015), we can virtually age the brain model and systematically evaluate the complex changes in the brain dynamics. This enables assessment whether the deterioration of certain cognitive processes or states are linked with the applied in-silico perturbations of the brain network models (Murphy et al., 2020; Pillai & Jirsa, 2017). If the generated functional patterns or states are then associated with a certain cognitive phenotype (Kelso, 2012; Pillai & Jirsa, 2017), the investigation of subject’s tethering between structural and functional connectivity via virtual aging allows understanding the brain’s adaptive capacities to react to individual cognitive decline (Naik et al., 2017).

We implemented the causal inference framework using biophysical modeling and resting-state functional magnetic resonance imaging (rs-fMRI) and diffusion-weighted imaging (DWI) data of 649 healthy participants of 1000BRAINS (Caspers et al., 2014). First, we analyzed the structural connectivity data and characterized its changes over age. Based on the major age-related changes, we systematically reproduced them in a subset of the youngest subjects’ brain models and virtually aged them to simulate a wide range of age-related and physiological parameters. After the results were benchmarked with the empirical data and simulation based on unperturbed DWI data, the parameters’ sweep of the modeling provides a landscape, in which brain fluidity changed as a function of age and global neuromodulation. We demonstrated that brain fluidity decreases when fiber tracts deteriorate and this leads to an increase in general global network neuromodulation as a potential scaffolding effect. Using deep learning networks, we independently confirmed the functional changes empirically in a large cohort (1000BRAINS) and assessed the most likely candidate mechanisms of healthy aging within the same causal framework of the Virtual Aging Brain (VAB). To demonstrate the causal relevance of the shift in network modulation in relationship to the Scaffolding theory of aging (Reuter-Lorenz & Park, 2014), we linked it to changes with age, sex and particularly cognitive performance, revealing further evidence for functional dedifferentiation and scaffolding in older adults.

## Results

We propose that the mechanistic inference framework of the VAB can causally link white-matter degeneration to age-related functional changes. By means of the computational framework of TVB, we designed virtual brains, i.e. data-driven brain network models integrating imaging data on SC to constrain network parameters, and we systematically generated a personalized ensemble of functional data (including FC and FCD variations) under various degrees of interhemispheric SC decrease. We specifically defined this process of artificial decline and assessment of the functional changes as “virtual aging”. Figure 1 summarizes the purposes and the steps of the VAB pipeline, which was meant to confirm the role of white matter degeneration in determining functional and cognitive changes during senescence as follows. Starting from various aging theories (Cabeza, 2002; Festini et al., 2018; Goh, 2011; Reuter-Lorenz & Park, 2014) that link the interindividual variability during senescence and the personal trajectories of age and cognition with each individual’s phenotype (comprised of structure and function, Figure 1.A), we designed individual brain-network models in the 1000BRAINS dataset in 2 scenarios. First, we estimated a model for each subject by constraining the brain network model parameters via the empirical connectome (*SC_emp_*) and we assessed whether we could generate realistic functional data (Figure 1.B). Second, we tested the process of “virtual aging” for the SCs of the 50 youngest subjects of 1000BRAINS by artificially decreasing their interhemispheric connections and hence generating a new set of connectomes (labeled as *SC*_α_, Figure 1.C). After verifying that the interhemispheric degeneration could drive age-related functional changes in both scenarios by comparing static and dynamic properties of FC with the empirical data (Figure 1.D), we eventually explored the tethering between structural connectivity and functional connectivity, expressed as the *G* modulation index, in relationship with age, the artificial decrease of SCs and cognition. Subsequently, we sought to test if the variability or trajectory of the SC-FC tethering, considered as one’s connectivity phenotype, might explain the heterogeneity of cognitive decline (Figure 1.E). Eventually, we wanted to independently validate the putative causal hypothesis of the VAB pipeline via Bayesian estimation of the tethering between SC and FC (Figure 1.F). Specifically, we retrieved the SC-FC link by means of a deep-learning approach, which merged a set of age-related functional changes from both the empirical and simulated data.

**Figure 1:**
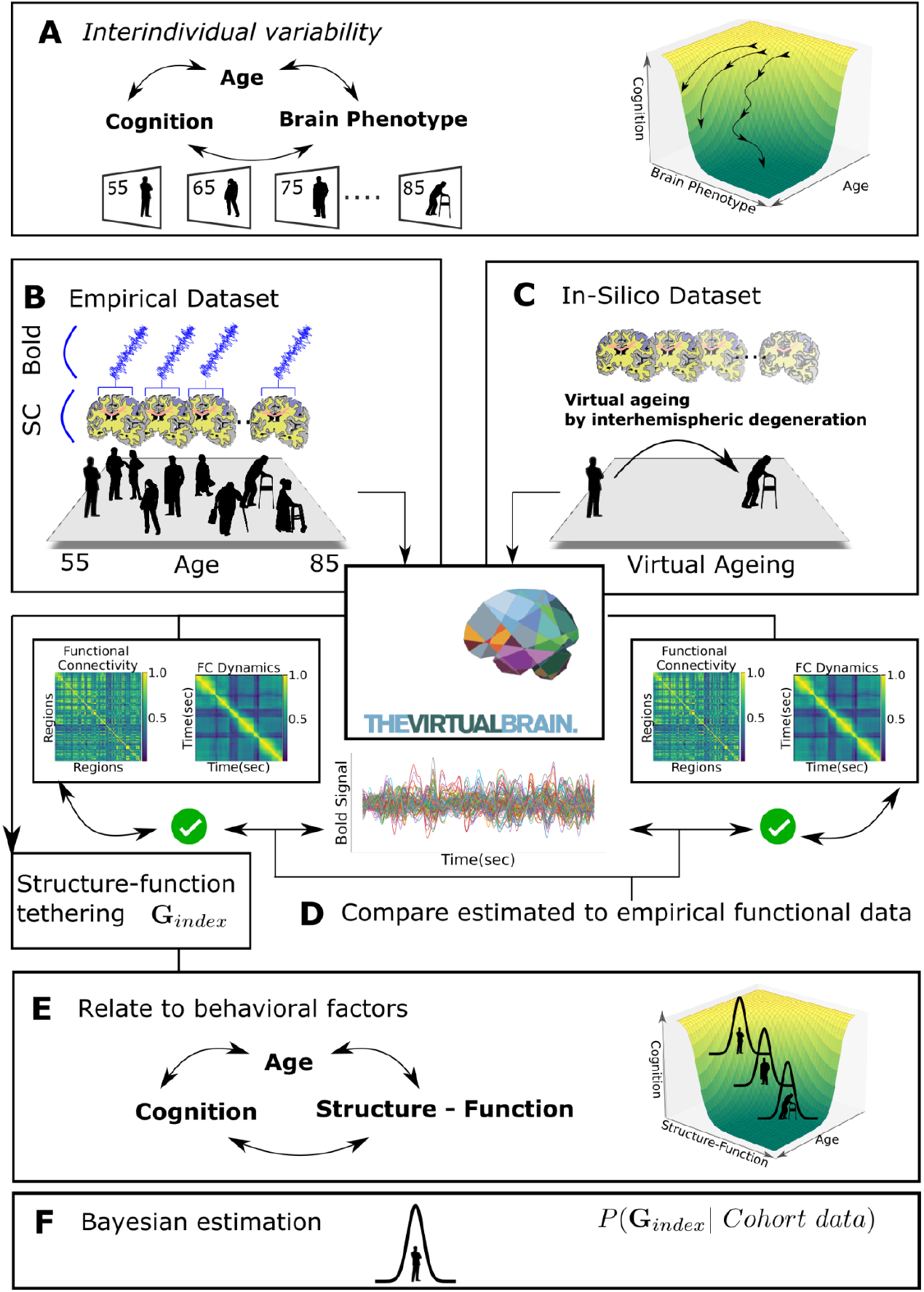
The mechanistic framework steps of the VAB to casually link wihte matter degeneration with age-related functional changes. (A) To explain the heterogeneity of cognitive performance during senescence, we designed personalized brain-network models in two scenarios. (B) First, we estimated virtual brains based on the empirical connectomes of 1000BRAINS dataset to generate functional data. (C) Second, we virtually age the brain models of the 50 youngest subjects structural connectomes by artificially declining the interhemispheric SC. (D) For both scenarios, we verified that the network models could realistically mimic in-vivo data by comparing the static and dynamical functional connectivity of the simulated data. (E) Thanks to the individual brain network models, we could derive an index of the causal link between structural connectivity and brain function, here labeled as *G* index to explain the heterogeneity of cognitive decline. (F) Eventually, we independently verified the estimation of our putative causal hypothesis using a Bayesian framework, which provided an uncertainty level of our estimation.

### R.1 Virtual aging results: the causal link between structural connectivity and functional connectivity and the relationship with cognition

From the DWI data of the 1000BRAINS participants (n = 649, age-range, mean: 67.2 year), we extracted individually weighted adjacency matrices for the brain network model (*SC_emp_*) using a brain parcellation of 100 regions (nodes) (Schaefer et al., 2018) (see details in Methods) and between which the collated white-matter tract densities (edges) are estimated. As suggested by previous work (Puxeddu et al., 2020), a preliminary investigation of *SC_emp_* revealed a strong age-related decline of interhemispheric fibers (Figure 2.C). To causally test whether white-matter degeneration could drive age-related functional changes, we designed brain network models as shown in Figure 2.A. Each model consists of a set of nodes or neural masses, which represent the mean field or the average activity of the associated brain regional neural population (El Boustani & Destexhe, 2009). The model chosen for each node is described in (Montbrió et al., 2015) and consists of two state variables that are bounded between an up-state of high activity and a down-state of fading activity. The nodes are connected via a graph, whose edges are represented by the SC weights. Effectively, the brain network has a dynamic for each region, which is influenced by the other regions thanks to the graph. To generate in-silico data, each brain network model was implemented in the open-source platform TVB, where the time-series for each region as well as the BOLD signals were simulated according to the Windkessel model (Friston et al., 2000) (Figure 2.A). Specifically, in-silico experiments were conducted in two scenarios, represented by the connectomes in Figure 2.B and 2.D. On the one hand, the brain network model was constrained by the empirical connectomes *SC_emp_* and we referred to the simulated data obtained with this model as age-simulation (also labeled as *sim*). On the other hand, we constrained brain network models via the connectomes of the 50 youngest subjects whose interhemispheric SC were homogeneously decreased from 0% to 100% by effectively masking the antidiagonal edges of the structural connectivity matrix (Figure 2.D). We referred to the functional data obtained under this variation of interhemispheric SC as α-simulation and virtually-aged data (also labeled as α − *sim*). To assure that our brain network models could reproduce age-related functional changes found in the empirical data, we calculated on both in-silico functional datasets (*sim* and α − *sim*) a set of summary statistics that represents the interhemispheric communication and its dynamical flexibility and compared them to those from the empirical functional data of 1000BRAINS. Since we suspected that SC deterioration had the greatest impact on the coactivation between hemispheres (Mollink et al., 2019; Roland et al., 2017) and we also wanted to investigate the impact of SC on brain fluidity (Courtiol et al., 2020; Hansen et al., 2015; Melozzi et al., 2019), we specifically computed the homotopic FC and the FCD variance difference. The homotopic FC is computed as the average of the FC of the same regions across hemisphere (e.g., prefrontal right and left), which are normally obtained as the diagonal of order 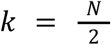 with N set to the total number of regions (Figure 2.E). The FCD variance is defined and was computed as the Frobenius norm of the *timeXtime* correlation matrix of the FC stream in Figure 2.E and the FCD variance difference is obtained as the difference between the interhemispheric FCD variance σ_*inter*_^2^ and the variance of full FCD σ_*full*_^2^ (Figure 2.E and see Methods). Additionally, each brain network model presents a set of free parameters that were tuned by sweeping the parameters in target range and selecting those that maximize σ_*full*_^2^ (see Methods). Among these, the global modulation *G* represents the overall influence of the SC-mediated input from the other regions on each node and we considered this parameter as an index of the modulation between structural and functional connectivity.

**Figure 2:**
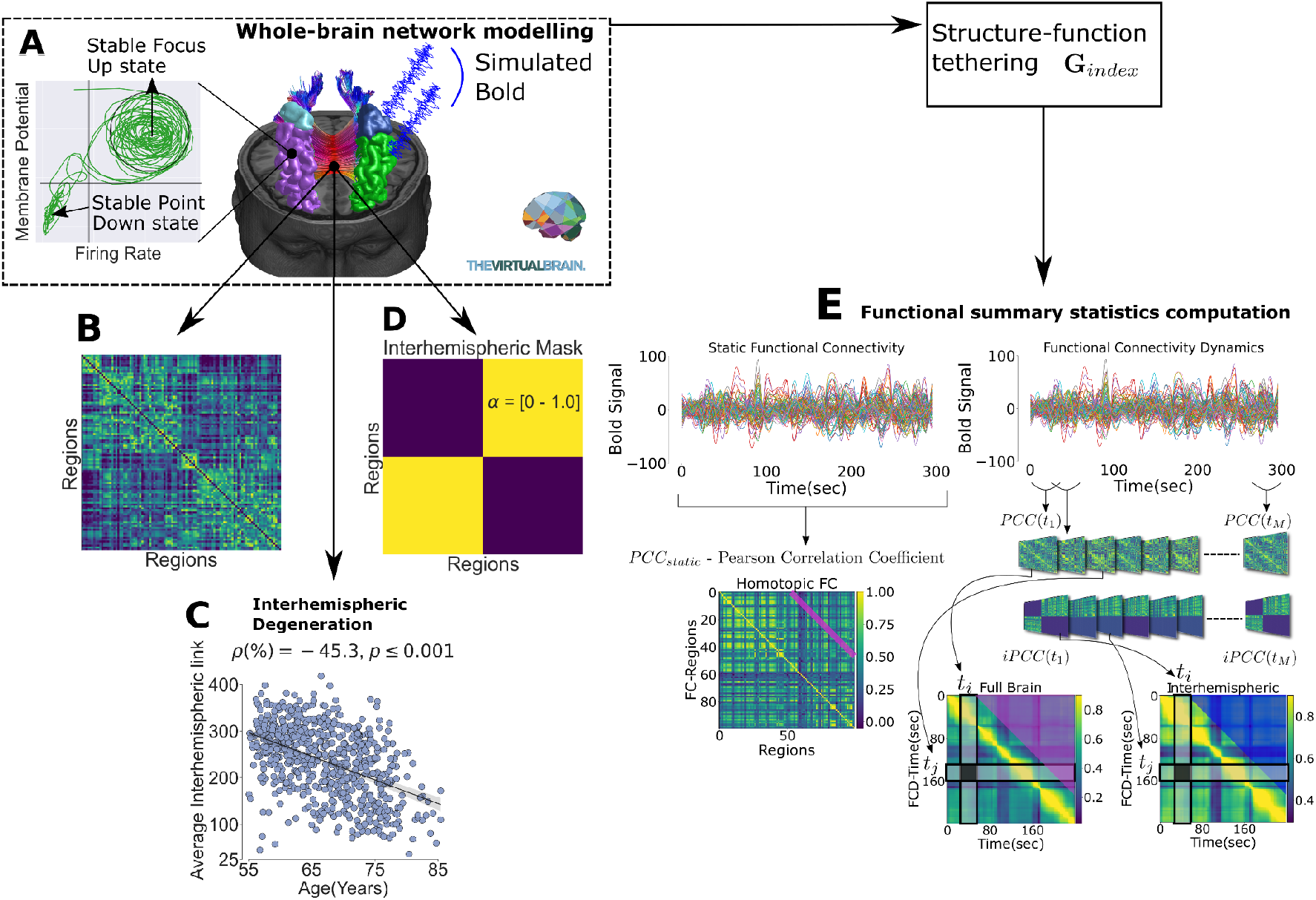
Detailed procedure of virtual aging to causally link white matter degeneration and age-related functional changes. (A) We designed virtual brains, i.e. brain network models composed of neural mass models as described in (Montbrió et al., 2015) and connected by the weights of the SC matrix in two scenarios. (B) First, we considered the empirical connectomes of 1000BRAINS. Second, based on the strong SC interhemispheric decline (C), we investigated artificial decrease of interhemispheric SC by masking the interhemispheric edges of the 50 youngest subjects (D) and testing various degrees of uniform changes (parameter α from 0% to 100%). By verifying that the simulated data could replicate the empirical functional changes, we defined the masking process as “virtual aging”. (E) We specifically test if we could replicate the decline of homotopic FC (magenta line in left panel) and the FCD variance difference, which is the difference of the variance of entries between the blue and the magenta triangle (right panel). See methods for details. Eventually, we could derive for each model a global parameter of modulation of SC onto the brain dynamics, which has been selected as the representative index of the causal tethering between SC and FC.

Figure 3 shows the results of the summary statistics as obtained for the empirical data, the simulation data and the virtually aged simulation data. In terms of homotopic FC, we found a negative age-trend for all three scenarios, the empirical data (Figure 3.A), the simulated data based on *SC_emp_* (Figure 3.B) and the simulated data based on *SC*_α_ (Figure 3.C) (Partial correlation corrected for sex and education: ρ_*emp*_ = 22.88% - *p* ≤ 0.001 - Figure 3.A, ρ_*sim*_ = −10.48% - *p* ≤ 0. 01 - Figure 3.B, Pearson Correlation - median(IQR): ρ_α−*sim*_ = −87% (14.8%) - Figure 3.C). Similar to the homotopic FC, we observed an age-related decline in the FCD variance difference (σ_*diff*_^2^) in all three datasets (Partial correlation corrected for sex and education: ρ_*emp*_ = −21.18% - *p* ≤ 0.001, - Figure 3.D, ρ_*sim*_ = −17.39% - *p* ≤ 0.001 - Figure 3.E, Pearson Correlation - median(IQR): ρ_α−*sim*_ =− 91%(6.6%) - *p* ≤ 0.001 - Figure 3.F). Panels 3.G and Figure 3.H show the sweep of the parameter *G* for each subject of the dataset *SC_emp_* and for each alpha connectome of the dataset *SC*_α_ for one target subject (out of 50, the remaining panels of *SC*_α_ can be found in the supplementary materials, Figure S3). Specifically, the FCD variance is shown as a heatmap for each considered *G* in the sweep procedure for each subject’s age (Figure 3.G) and for each alpha of a virtually aged subject (Figure 3.H). They both suggest an increase of the *G* associated to the maximum of the FCD variance. Since *G* is by definition representative of the impact of SC over the brain’s functional architecture in the brain network model, we decided not only to address whether *G* was related to age and the artificial interhemispheric decrease, but if it could also be associated with the heterogeneity of cognitive decline.

**Figure 3:**
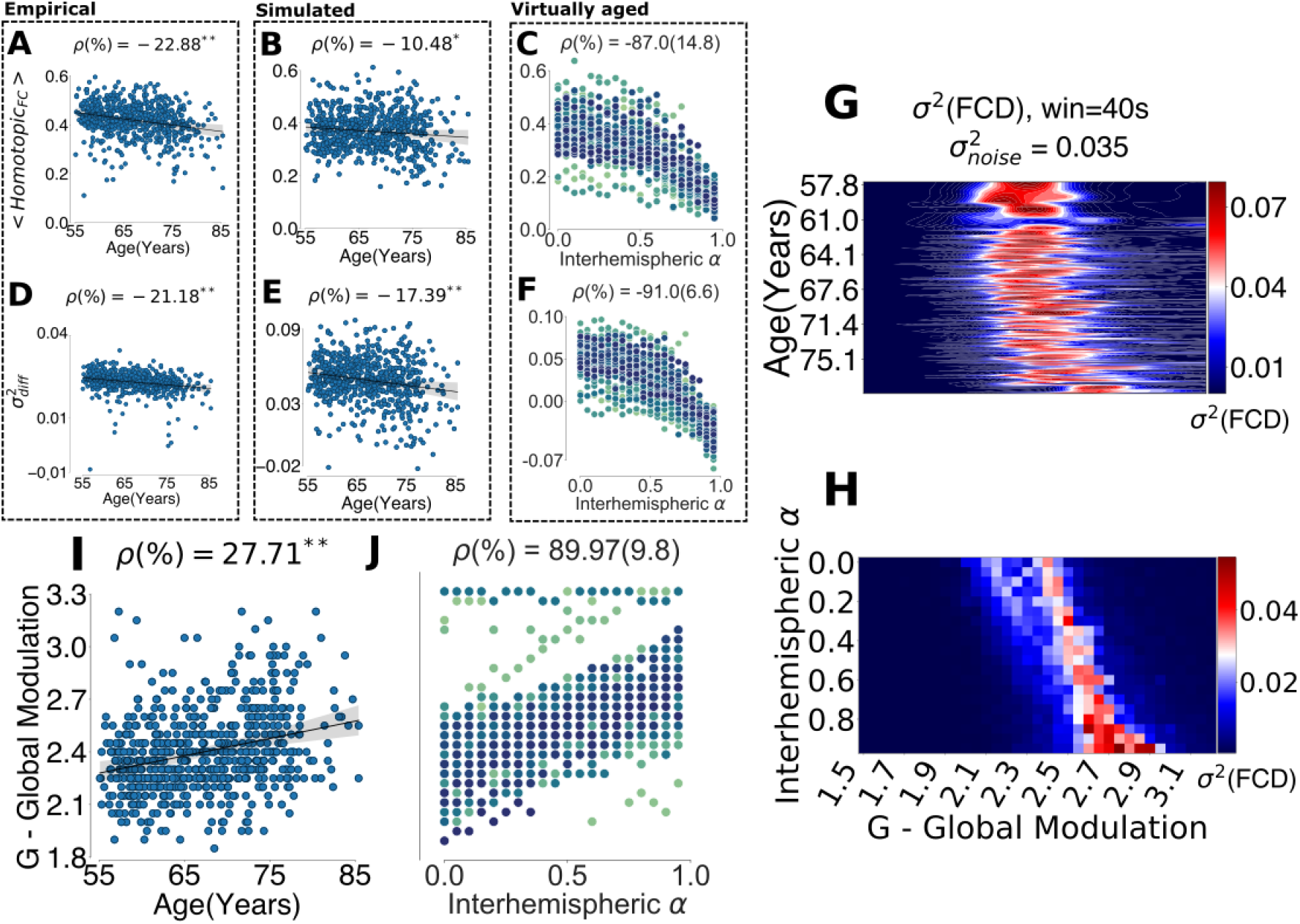
(A-B-C) The top three panels show the trend for homotopic FC for the empirical, the simulated and the virtually aged data in relation to age or α. Specifically, the first two columns report the trend for each subject in function of age, while the latter shows the trend for the 50 youngest subjects virtually aged via interhemispheric αparameter. We observed negative trend for all three cases (** *p* ≤ 0.001, * *p* ≤ 0.01). (E-D-F) Similarly to the homotopic FC, we reported the in the FCD variance difference (σ_*diff*_^2^) in all three datasets and we also observed a negative trend for all three cases. (G) Heatmap the simulated FCD variance σ_*full*_^2^ for the entire 1000BRAINS dataset along two dimensions: the *G* modulation index (range [1.5 - 3.2]) and the age of each connectome. (H) Heatmap the simulated FCD variance σ_*full*_^2^ for a virtually-aged target subject along two dimensions: the *G* modulation index (range [1.5 - 3.2]) and the interhemispheric α (see the remaining heatmaps in Figure S3). (I-J) The G associated to maximum of FCD variance, as index of the SC-FC tethering, shows a increasing G with age (Partial correlation corrected for sex and education) and interhemispheric decrease α (Pearson Correlation - median(IQR)).

To systematically address this question, the *G* modulation index was related to age, sex and cognitive performance. Results revealed both, age as well as sex to be significantly related to the *G* modulation index (ANCOVA corrected for education, sex: *F* = 20.13, *p* ≤ 0.001, η^2^ = 0. 03, age: *F* = 53.16, *p* ≤0.001, η^2^ = 0.075). Thereby, increasing age was associated with a stronger linkage between SC and FC (Partial Correlation, corrected for education and sex: ρ = 27.71%, *p* ≤ 0.001, Figure 3.I). Importantly, testing the association between the *G* modulation index of both *SC_emp_* and *SC*_α_ revealed similar results indicating a robust increase of the *G* modulation index not only as a function of age but also as a function of α (Pearson Correlation - median(IQR): ρ = 89.97%(9.8%), Figure 3.J), at this demonstrating the ability of the model to reflect a relevant aging-associated phenomenon. For sake of completeness, the other free parameter of network model, the noise variance σ^2^ (see Methods for further explanation), showed a flat trajectory centered on σ^2^ = 0.035 and σ^2^ = 0.04 in the function of age (Partial correlation corrected for sex and education: ρ = 1.42%, *p* = 0. 71, Figure S1.A in the supplements) as well as in the function of α (Pearson Correlation: ρ = 5. 87%(43. 7%), Figure S1.B in the supplements).

With regards to sex, females were found to exhibit significantly lower SC modulation on the FC as compared to males. These sex-related differences were constant across age as both sexes showed comparable correlations between the *G* modulation index and age, i.e. with males showing a constantly higher tethering between SC and FC (Partial Correlation, corrected for education, women: ρ = 27.02 %, *p* ≤ 0.001; men: ρ = 27.79%, *p* ≤ 0.001; Fisher’s Z: *p* = 0.46, Figure 4.A). With regards to aging theories indicating that changes in the interplay between structure, function, and connectivity may explain the variability of cognitive performance during the aging process (Festini et al., 2018; Reuter-Lorenz & Park, 2014), we tested whether the variability of cognitive performance could indeed be explained by the *G* modulation index obtained with *SC_emp_* using ordinary least square regression (corrected for confounding effects of age, sex and education). All associations between cognitive performance and the SC-FC linkage can be found in Supplementary Table 1. We found the *G* modulation index to be significantly related to the performance in concept shifting (β_*G*_ = −19.94, *p_G_* = 0.043, β_*age*_ =− 1.54, *p_age_* ≤ 0.001, β_*sex*_ = 3.072, *p_sex_* = *n.s*., β_*edu*_ = 3.93, *p_edu_* ≤ 0.001, Bonferroni corrected, Supplementary Table 1). Here, a higher G-coupling index was found to be associated with lower performance. Splitting the group by the performance median (−45.235) into low performing (LP, *n* = 327) and high performing subjects (HP, *n* = 322), we interrogated whether performance strength may be associated with different age-trajectories of the *G* index. Indeed, we found low performers to not only be associated with a significantly higher *G* modulation index as compared to high performers (ANCOVA corrected for age, sex and education, HP/LP-CSH: *F* = 5.5, *p* = 0.019, η^2^ = 0.008), but also to have a stronger age-related increase of the *G* index (Partial Correlation corrected for sex and education, LP: ρ_*LP*_ = 31.40 %, *p_LP_* ≤ 0.001; HP: ρ_*HP*_ = 18.96 %, *p_HP_* ≤ 0.001; Fisher’s-Z: *p* = 0.046, Figure 4.B). As the age-related differences of the *G* modulation index revealed a non-linear trend (see Figure 3.G-3.I), we additionally split the whole group into older and younger participants (cut by median = 67 years). Thereby, the older group was found to show significantly different age-related changes in the *G* modulation index with low performers showing a stronger increase of the linkage between structural connectivity and functional connectivity across ageing (Partial Correlation corrected for sex and education: LP: ρ_*LP*_ = 36.30 %, *p_LP_* ≤ 0.001; HP: ρ_*HP*_ = 5.66%, *p_HP_* = 0.48; Fisher’s-Z: *p* = 0.002, Figure 4.D), while the younger low performers did not present a different trajectory from the high performing group (Partial Correlation corrected for sex and education: LP: ρ_*LP*_ = 12.49 %, *p_LP_* = 0.115; HP: ρ_*HP*_ = 5.49%, *p_HP_* = 0.49; Fisher’s-Z: *p* = 0.264, Figure 4.C). Notably, we additionally found the global G-coupling index to be significantly related to verbal memory (β_*G*_ =− 4.94, *p_G_* = 0.044, β_*age*_ =− 0.457, *p_age_* ≤ 0.001, β_*sex*_ = 6.376, *p_sex_* ≤ 0.001, β_*edu*_ =1.11, *p_edu_* ≤ 0.001, Bonferroni corrected, Supplementary Table 1). This, however, was not accompanied by a stronger correlation between performance and the *G* modulation index in low performers, neither in the older age - group nor in the entire cohort.

**Figure 4:**
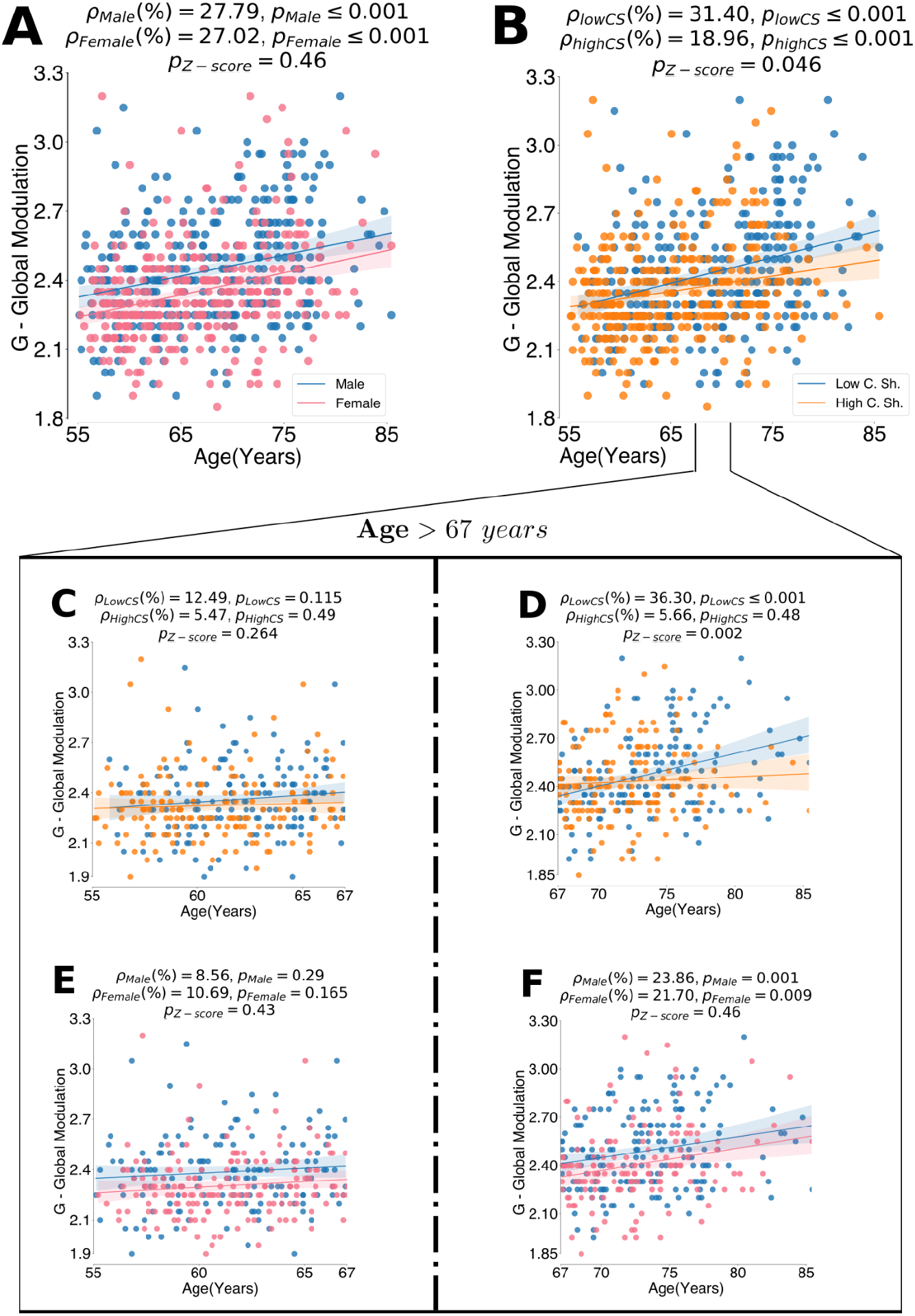
(A) The trend of global modulation G split by sex showed that women (pink) and men (blue) have similar increase in terms of G trend with age, as supported by the Fisher’s Z (B) The global modulation G trend during ageing split by concept shifting show a steeper G increase for the low performers (blue) than high performers (orange). (C-D) The same trend in (B) split by the median age 67 years showed different trends for the younger subjects and older ones. The former shows no significant trend with age and no difference in terms of cognitive scores, while the older patients show a more prominent difference between the two groups than the entire cohort. (E-F) The same trend in (A) split by the median age 67 years for the younger subjects and older ones. Both in younger and in the older cohort, the age-trend of G presents similar patterns between men and women.

### R.2 The Bayesian validation: Simulation-based Inference

To ensure the validity of the virtual aging mechanistic framework, we independently confirmed the causal link and the increase of the structural connectivity-functional connectivity tethering by means of the simulation-based inference (SBI) framework. Unlike the parameter sweep approach, this Bayesian framework used a sequential neural posterior estimation (SNPE) to obtain a posterior estimation of the modulation index *p*(*G_SBI_*|*X_empirical_*), based on a set of summary statistics of the empirical data (*X_empirical_*) Utilizing a deep learning approach (see Methods), SBI estimates the likelihood function of the Bayes rule for each subject by using a budget of simulated data (2000 simulations) and a set of informative features related to functional changes with age, such as homotopic FC, the FCD variance difference and the standard deviation of the interhemispheric FC stream (further details in the Methods section). Given the uniform prior distribution on the parameter *G* and the trained deep-learning network on the simulated data representing the likelihood function, we directly derived the posterior distribution of the parameter *G* by employing the neural density estimator, using the same summary features obtained from the empirical data. In order to provide a single value estimation for each subject, we considered the mean of the posterior distribution as the estimated *G* (see Methods for more details). By applying SBI, we observed that the obtained global *G* modulation increased with age (ρ = 25.38%, *p* ≤ 0.001, Figure 5.A). Moreover, we also observed an agreement between the *G* estimated with parameter sweep and that obtained with SBI (ρ = 41.3%, *p* ≤ 0.001, Figure 5.B), which independently confirmed the observation of the causal link between the two domains obtained with the parameter sweep and the age-related increase of the SC-FC tethering. We also verified the posterior distribution for each subject to confirm that the overall mean trend of *G* meaningfully increased with senescence (Figure 5.C and 5.D). Moreover, we estimated the global *G* used to generate data for a target connectome to validate that the SBI approach indeed encompassed the ground truth of the in-silico simulated data for a subject, (see Figure S2, in supplementary materials).

**Figure 5:**
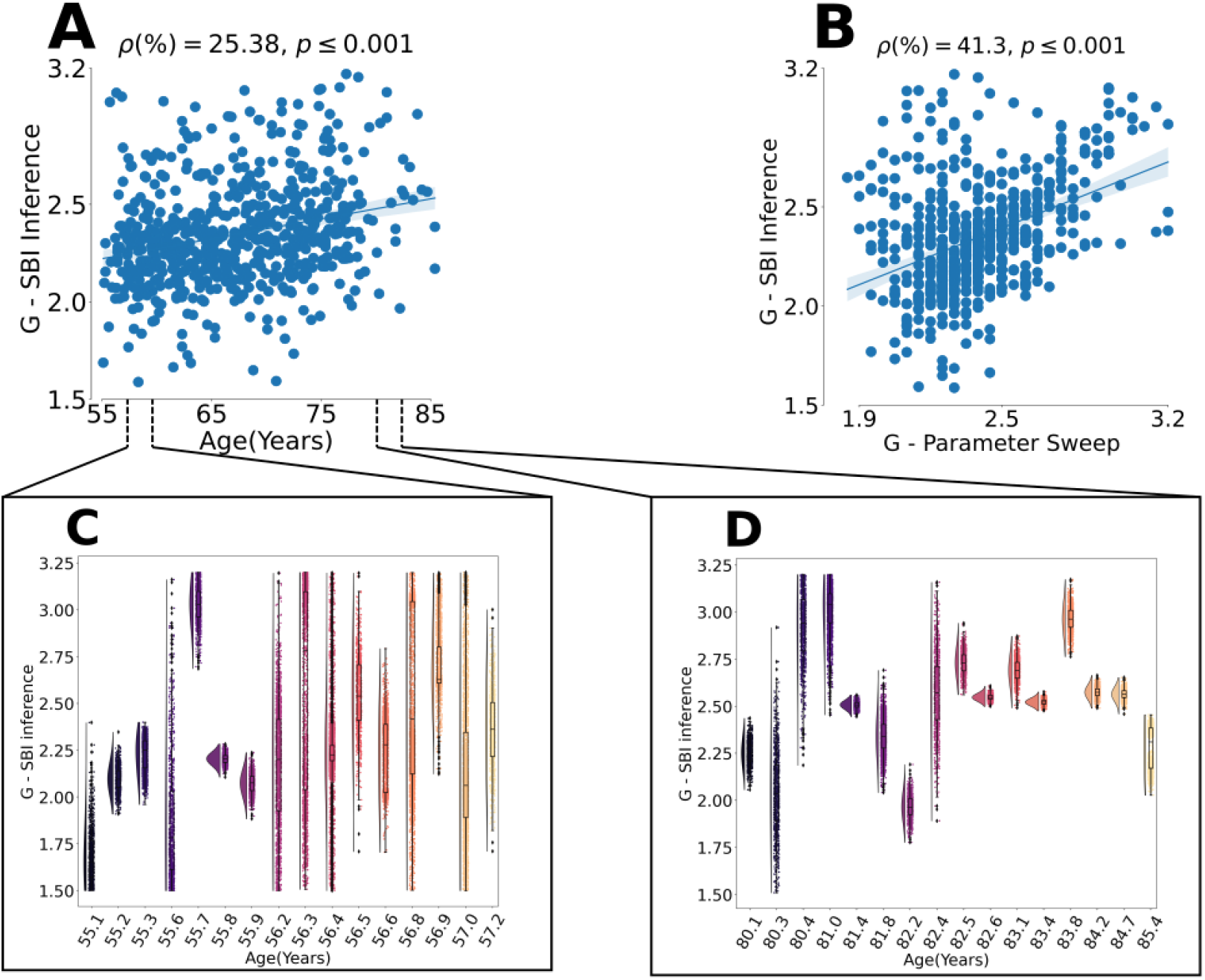
(A) The global modulation *G* obtained by SBI framework (see Methods) also increased with age similarly to Figure 3.I. (B) The Bayesian approach validated the causal link estimation since the *G* modulation index obtained with SBI showed agreement with the G obtained by parameter sweep. (C) The posterior distribution of G obtained with SBI for the subjects in the lifespan [55-57] years. (D) The posterior distribution of G obtained with SBI for the subjects in the lifespan [80-85] years.

## Discussion

The here established causal inference framework disentangles several coexisting mechanisms, namely the interhemispheric deterioration and the cognitive decline, during the healthy aging process of the brain. Building virtually aged brain network models, additionally confirmed by deep learning networks, we empirically replicated the age-related functional changes in a large cohort (1000BRAINS) and demonstrated that with increasing age the network global neuromodulation of the brain increases. This was achieved by manipulating interhemispheric SC as the major influencing factor in terms of both, static as well as dynamic functional connectivity. We operationalized causal structural connectivity changes by a novel masking approach, in which the degree of fiber deterioration is implemented as weight in each individual connectivity matrix for virtual aging. When we modeled the progression of brain dynamics across age, we took into consideration the coexisting changes of SC and global network modulation and showed that deterioration of interhemispheric SC leads to a loss of fluidity in brain dynamics, which resulted in an increase in global network modulation. Our independent empirical finding demonstrates that the brain might undergo a process of dedifferentiation and consequent scaffolding for the gradual deterioration of structural connectivity during senescence, especially in older adults with poor cognitive performance, as suggested by the Scaffolding theory of aging.

Existing neurocognitive theories of aging have argued that functional changes following structural deterioration are either beneficial or detrimental. Despite numerous empirical investigations, there is no clear consensus on the mechanisms underlying the dynamic processes that occur over the lifespan, probably largely due to the presence of coexisting mechanisms. In line with previous studies focusing on age-related SC differences (Puxeddu et al., 2020; Zhao et al., 2015), the interhemispheric SC is the most prominent structural factor showing the strongest age-related decreases. This anatomical driver was used to virtually reproduce an age trajectory for each individual by masking interhemispheric structural connections. Applying the computational models to both, the empirical *SC_emp_* dataset as well as the virtually reproduced *SC*_α_ datasets revealed decreases in the homotopic FC as well as lower FCD for all simulated functional data. The functional connectivity between homotopic regions has been previously reported to be highly age-characteristic (Chan et al., 2014; Senden et al., 2017; Tzourio-Mazoyer et al., 2016; Zuo et al., 2010) including the dynamical interplay between the two hemispheres (Escrichs et al., 2021; Xia et al., 2019). Our findings showed that the modeling framework of the Virtual Aging Brain (VAB) can bridge the explanatory gap between the structural connectivity changes and the brain’s functional reorganization during the aging process. With the current results we can, for the first time, mechanistically confirm the previous assumptions that inter-hemispheric SC serves as a pivotal basis for homotopic FC (Jin et al., 2020; Mollink et al., 2019; Roland et al., 2017). Specifically, our results show age-related decreases of not only homotopic FC but also interhemispheric FCD to be driven by inter-hemispheric SC decreases, suggesting an emergence of functional dedifferentiation in older adults (Chan et al., 2014; Reuter-Lorenz & Park, 2014).

Thus, our approach goes beyond existing neurocognitive theories of aging, which provide arguments based on interpretation of empirical observations. The Compensation related utilization of neural circuit hypothesis (CRUNCH theory) proposes that overactivation is a common response across all age groups when cognitive demand is high (Festini et al., 2018). Similar observations are made with regard to hemispheric asymmetry reduction in older adults (HAROLD theory) but emphasize the overactivity exhibited by task-unrelated contralateral brain regions in high task-demand conditions (Cabeza, 2002). The ‘scaffolding theory of aging and cognition’ (STAC) proposes positive plasticity or scaffolding, which might induce functional dedifferentiation with age by enacting bilateral recruitment or invoking primary networks to maintain behavioral integrity (Reuter-Lorenz & Park, 2014). These subsequent functional changes reflect an adaptive recalibration process to maintain functionality despite the decline of the underlying white matter pathways (Reuter-Lorenz, 2014). However, this recalibration only happens if there are no other external factors that could preserve the neural substrate (such as education or training). The essential observation in these theories is that the brain must adopt alternative pathways to preserve information exchange (Naik et al., 2017), potentially resulting in an over-recruitment of intermediate brain regions passing on information, to limit cognitive decline. This phenomenon of functional dedifferentiation might occur as a reaction to SC degeneration (and, more in general, the neural substrate), which is supposed to be the primary cause of the poor cognitive performance. However, none of the empirical accounts offers a causal inference perspective and, thus, does not progress beyond hypothesis. Our findings now overcome this and demonstrate that a constantly higher recruitment of brain regions results in lower functional diversity and similar brain activation patterns between hemispheres (Lou et al., 2019), i.e. decreasing variance of functional dynamics indicating a lower interhemispheric metastability. Given a higher metastability to reflect the brain’s ability to dynamically transition between different cognitive states (Power et al., 2011), lower fluidity (as captured by FCD) can be understood as a reduced speed of mind state reconfigurations in response to external influences (Lee et al., 2019; Xia et al., 2019).

In line with previous work indicating the SC-FC linkage to become stronger across aging (Zimmermann et al., 2016), we found increasing age to be characterized by a higher global network modulation, which can be interpreted as an indicator for how much a structural connection modulates the existence of a functional connection (Fukushima et al., 2018). This could hint at the fact that by the increasing need for alternative pathways that compensate for damages of direct structural links, an overall higher proportion of the intact structural connectome is utilized (also reducing the capacity of flexible functional reconfigurations). In fact, FCD decreases are suggested to reflect the neurological counterpart of the processing speed theory, implying decreases in the metastability to reduce the speed of information processing which then accounts for overall slower behavioral responses (Finkel et al., 2007; Salthouse, 1996; Xia et al., 2019). Remarkably, in the current work we indeed found that lower levels of in-silico FCD difference is lower in older adults (functional dedifferentiation in STAC theory terms), while the tethering between SC and FC is higher, especially in poor cognitive performers (Figure 4). This may suggest that the individual modeling amplifies this reduced brain fluidity for the subjects with a lower performance in concept shifting, suggesting a scaffolding effect as expressed by the STAC theory.

As discussed by various authors (Goh, 2011; Naik et al., 2017; Reuter-Lorenz & Park, 2014), the linkage between structural connectivity and functional connectivity is suggested to be a cornerstone in explaining the cognitive decline and its associated interindividual variability. In addition to the SC-FC interaction changes with age, we were able to demonstrate that the global network modulation is negatively related to verbal memory and concept shifting (Supplementary Table 1), which was particularly driven by the older subjects in the here assessed 1000BRAINS cohort. Moreover, splitting the group into low and high performers revealed the low performing group to be depicted by a faster increase of the global modulation parameter *G* supporting the notion that the tethering between SC and FC is indeed highly important for the progression of cognitive performance (Figure 4). Collectively, our findings indicate a greater deterioration of the interhemispheric structural connectivity to increase the modulation by the SC adjacency matrix on the brain functional dynamics as potential scaffolding reorganization of the brain. This tighter interaction generated dedifferentiated functional aging patterns, i.e. homotopic FC and FCD decreases, that ultimately led to cognitive decline. Remarkably, the scaffolding effect of the *G* modulation seems not to happen for the high performing group, which suggests a process of brain maintenance occuring for other intervening factors (Reuter-Lorenz & Park, 2014).

There have been various attempts in the literature trying to explain the complex relationship between SC and FC (Fukushima et al., 2018; Suárez et al., 2020; Zimmermann et al., 2016, 2019), but they all relied on data-driven and statistical methods, which limited the interaction between the two modalities to percolation (Saggio et al., 2016; Wein et al., 2021). Straathof et al. (Straathof et al., 2019) pointed out that the estimation of the SC-FC relationship may be characterized by a large unexplained variability possibly due the limited information of the regional averaged FC. Although data-driven approaches showed strong variations in the SC topology leading to high variations in the overall static FC (Roland et al., 2017; Wendelken et al., 2017; Zimmermann et al., 2016), the mechanistic framework of the Virtual Aging Brain (VAB) introduced in the current work provides a brain-network model that did not limit the relationship between SC and FC to only the static properties of FC, but extends it to its dynamic properties (FCD). Hence, it could also track the evolution of this complex interaction during senescence. Thanks to the capacity to handle the dynamic and static properties of FC and the possibility of a nonlinear decline of artificial interhemispheric functional data, the VAB approach underpins a nonlinear relationship between SC and FC, which comes closer to linear or 1:1 modulation of structure over function with age and cognitive decline.

To validate whether the brain network models could actually estimate the relationship between structure and function, we applied a Bayesian framework (SBI) to infer the full posterior of parameters from the joint distribution of empirical functional data and a low-dimensional set of summary statistics (Cranmer et al., 2020; Gonçalves et al., 2020; Hashemi et al., 2020). By considering the same premises on the metastable behavior of the brain, we employed a simulation-based inference that retrieves model parameters with the same age-declining FC and FCD features. Unlike the parameter-sweep, the SBI could rely on the individual estimate of global modulation by both considering the SC and functional data in a low dimensional space. Therefore, the Bayesian approach independently provides a ground truth value of change of global modulation, which becomes interpretable through the mechanistic framework virtual aging, in which we separate coexisting influences of global modulation and structural deterioration. Importantly, we systematically demonstrate the direction of the scaffolding and dedifferentiation process, which always was directed towards increasing global modulation. Our Virtual-Aging framework is not only capable of estimating the interaction between structural connectivity and functional connectivity on an individual basis, but it can determine the uncertainty of the tethering at subject level.

## Conclusions

We have demonstrated that functional brain changes are causally linked to age-related white matter degeneration. Specifically, inter-hemispheric SC decline modifies brain function in the sense that older adults are characterized by a reduced fluidity in functional brain dynamics as measured by lower FCD and a lower interhemispheric metastability. Lower FCD being understood as a reduced capacity of mind state reconfigurations, the brain’s ability to dynamically switch between cognitive states may be reduced. This could indeed explain the decline in the performance of concept shifting in older adults who showed a higher SC-FC modulation. As SC causally modifies the brain’s functional capacity to switch between brain states, it manifests behaviorally in a performance decline. This interpretation is further corroborated by our independent demonstration that individual functional brain signals are better explained by increased values of scaled SC under a causal brain network hypothesis, suggesting a scaffolding effect linked to the reduced brain fluidity. To our knowledge, this is the first mechanistic evidence of scaffolding linked to functional dedifferentiation in aging leading to cognitive decline demonstrated within a subject-specific causal inference framework in a large cohort.

In summary, the present model does not only capture how age-related SC changes impact the homotopic FC during the aging process, but additionally shows that the effects on brain’s fluidity or FC dynamics and its associated decline in the later stages of life can be modeled. Importantly, the intrinsic nonlinearity of the brain network model suggests that SC-FC tethering changes with age cannot be mapped in a one-to-one fashion. Based on our findings, we can assume a greater deterioration of the interhemispheric structural tracts to increase the modulation by the SC adjacency matrix on the brain dynamics. Assuming an optimal tuning state of each brain, our model demonstrated a shift of this sweet spot towards higher values, reflecting greater global network modulation. The tighter interaction between structural connectivity and functional connectivity generated dedifferentiated functional aging patterns, i.e. decreases in brain fluidity, which ultimately led to cognitive decline.

## Acknowledgments

This project was partially funded by the German National Cohort and the 1000BRAINS-Study of the Institute of Neuroscience and Medicine, Research Centre Jülich, Germany. We thank the Heinz Nixdorf Foundation (Germany) for the generous support of the Heinz Nixdorf Study. We thank the investigative group and the study staff of the Heinz Nixdorf Recall Study and 1000BRAINS. ML would like to thank Paul Triebkorn for the Visualization support and both Viktor Sip and Lionel Kusch for their insights on programming with HPC facilities. This project/research has received funding from the European Union’s Horizon 2020 Framework Programme for Research and Innovation under the Specific Grant Agreement No. 945539 (Human Brain Project SGA3). The authors also wish to acknowledge the financial support of the following agencies: the French National Research Agency (ANR) as part of the second “Investissements d’Avenir” program (ANR-17-RHUS-0004, EPINOV), European Union’s Horizon 2020 research and innovation programme under grant agreement No. 785907 (SGA2) Human Brain Project, PHRC-I 2013 EPISODIUM (grant number 2014–27), the Fondation pour la Recherche Medicale (DIC20161236442), Virtual-BrainCloud (grant number 826421), the SATT Sud-Est (827-SA-16-UAM) for providing funding for this research project, the Initiative and Networking Fund of the Helmholtz Association (SC), the Psychiatric Imaging Network Germany (PING) project (BMBF 01 EE1405C).

## Materials and Methods

### M.1 Sample

Participants of the current study are based on the 1000BRAINS project (*n* = 1314, 18-87 years, 582 females) (Caspers et al., 2014), a population-based study which was designed to analyze the normal aging process, i.e. the variability of brain structure, brain function and its relation to behavioral, environmental and genetic factors. 1000BRAINS is a subsample of the 10-year follow-up cohort of the Heinz Nixdorf Recall Study, an epidemiological population-based study that investigates risk factors for atherosclerosis, cardiovascular disease, cardiac infarction, and cardiac death (Schmermund et al., 2002), as well as the associated Multi-Generation Study. For 1000BRAINS, no exclusion criteria other than eligibility for MRI measurements (Caspers et al., 2014) were applied as the project aims to characterize aging at the general population level. Eligibility for MRI measurements included any history of neurosurgery, cardiac pacemakers, coronary artery stents, surgical implants or prostheses in head or trunk, tattoos or permanent make-up on the head. Participants gave written informed consent prior to inclusion in 1000BRAINS and MR imaging was only conducted as there were no dental implants found which could cause artefacts in the brain images and as participants did not experience claustrophobia. The study protocol of 1000BRAINS was approved by the Ethics Committee of the University of Essen, Germany. In the current work we focused on particularly older adults (above 55 years, *n* = 970), from which a total of 871 had functional scans available. 16 participants had to be excluded due to insufficient data quality. Of these 855 subjects, 718 also had diffusion data available, from which 649 were of good quality (see more details under Imaging). The current sample therewith comprises a total of *n* = 649 participants (age-range = [51.1-85.4] years, mean age = 67.2 years, *n_females_* = 317).

To test how the SC-FC tethering and the causal link between white matter degeneration and functional changes could explain to interindividual variability of cognitive decline, we took advantage of the comprehensive neuropsychological assessment that was conducted within 1000BRAINS and included 16 cognitive performance tests into the current study (see Supplementary Table 2, for more details also see (Caspers et al., 2014; Jockwitz et al., 2017; Stumme et al., 2020)).

### M.2 Imaging

All participants were scanned using a 3T Siemens Tim-TRIO MR scanner (32-channel head coil, Erlangen, Germany) located at the Forschungszentrum Jülich in Germany. Different imaging sequences, i.e. anatomical, diffusion and resting-state, were used in the current study to analyze the structural and functional connectivity tethering (for detailed description of the 1000BRAINS study protocol also see (Caspers et al., 2014)). For the anatomical image, a 3D high-resolution T1-weighted magnetization-prepared rapid acquisition gradient-echo (MPRAGE) anatomical scan (176 slices, slice thickness 1 mm, repetition time (TR) = 2250 ms, echo time (TE) = 3.03 ms, field of view (FoV) = 256 × 256 mm2, flip angle = 9°, voxel resolution 1 × 1 × 1 mm3) was acquired to perform a surface reconstruction. For structural connectivity, diffusion tensor images were acquired using the following parameters: a HARDI protocol with a subset (60 directions) EPI with TR = 6.3s, TE = 81ms, 7 b0-images (interleaved), producing 60 images with b = 1000s/mm2 and a voxel resolution of 2.4×2.4×2.4mm3; and a second HARDI subset (120 directions) EPI with TR = 8s, TE = 112ms, 13 b0-images (interleaved), producing 120 images with b = 2700s/mm2 and a voxel resolution of 2.4×2.4×2.4mm3. And lastly, for resting-state functional connectivity, we used a blood-oxygen level dependent (BOLD) gradient-echo planar imaging (EPI) sequence with 36 transversally oriented slices (slice thickness 3.1 mm, TR = 2200 msec, TE = 30 msec, FoV = 200 × 200 mm2, voxel resolution 3.1 × 3.1 × 3.1 mm3), lasting for ~11 minutes and producing 300 volumes. Of note, during resting-state image acquisition, participants were instructed to keep their eyes closed, be relaxed, let their mind wander and not fall asleep, which was assured by post-scan debriefing.

#### M.2.1 Structural Image Processing

Using the CAT12 toolbox implemented in SPM12 (Ashburner, 2009), we created tissue probability maps (TPM) for grey matter (GM), white matter (WM) as well as corticospinal fluid (CSF) from the participant’s T1 data. These brain masks were used to optimally extract the brain from the T1 data by superimposing and thresholding them at 0.5 (small enclosed holes were filled). The T1 brain was bias field corrected, rigidly aligned to MNI152 template space and resampled to 1.25 mm isotropic voxel size. We corrected the diffusion MRI data for eddy current and motion artifacts including interpolation of slices with signal dropouts (Andersson et al., 2016; Andersson & Sotiropoulos, 2016). Subsequently, all diffusion data were visually controlled for ghosting, remaining signal dropouts or very noisy data and volumes or datasets that were considered as suboptimal were excluded from further analysis (*n* =− 69). For dMRI - T1 alignment, the first b0 images from each dMRI data with b1000 and b2700 were extracted and rigidly aligned to T1 dataset using mutual information as cost function (Wells et al., 1996). Based on the estimated transforms, the dMRI data were transferred to the individual T1 space, separately for both b-values. Implicitly, the realignment resamples the data to 1.25mm and rotates the b-vectors according to the corresponding transformations. As there are no field maps or b0 volumes with opposite EPI readout directions available in the current study, we computed Anisotropic Power Maps (APM) from the b2700 dMRI data in 1.25 mm space to account for susceptibility artifacts and optimize image registration (Dell’Acqua et al., 2014). APM contrasts are very similar to those of the T1 image and therefore provide an optimized base for image registration. Hence, APMs were used to compute the non-linear transformation from diffusion to anatomical space, thereby taking EPI induced distortions into account. These non-linear transformations were then used to transform the TPMs to diffusion space. Finally, the two datasets with b1000 and b2700 were merged into one single file and corrected for different echo times. This correction was computed by a voxelwise multiplication of the b2700 data with the ratio of the non-diffusion-weighted data respectively for the two datasets. Lastly, local modeling and probabilistic streamline tractography were performed using the MRtrix software package version 0.3.15 (Tournier et al., 2012). We computed the constrained spherical deconvolution (CSD) local model using multi-tissue CSD of multi-shell data (Jeurissen et al., 2014) with all shells and a maximal spherical harmonic order of 8. Ten million streamlines were computed with dynamic seeding in the gray-white matter interface for every subject using the probabilistic iFOD2 algorithm with a maximal length of 250 mm and a cut-off value at 0.06.

#### M.2.2 Functional image processing

For each participant, the first four echo-planar imaging (EPI) volumes were discarded. To correct for head movement, a two-pass procedure was performed using affine registration: first, aligning all functional volumes to the participant’s first image and second, to the mean image. Functional images were then spatially normalized to the MNI152 template (Holmes et al., 1998) using the “unified segmentation” approach by (Ashburner & Friston, 2005). This was preferred to normalization based on T1 weighted images as previous studies indicated increased registration accuracies (Calhoun et al., 2017; Dohmatob et al., 2018). Furthermore, to identify and remove motion-related independent components from functional MRI data, we applied the data-driven method ICA-based Automatic Removal Of Motion Artifacts [ICA-AROMA (Pruim et al., 2015)]. According to previous suggestions indicating a combination of AROMA and global signal regression to minimize the relation between motion and resting-state Functional Connectivity (FC) (Burgess et al., 2016; Ciric et al., 2017; Parkes et al., 2018), we additionally performed global signal regression in the current study. Lastly, all rs-fMRI images were bandpass filtered (0.01 – 0.1 Hz). To assure data quality, we performed the established algorithm by (Afyouni & Nichols, 2018) on the preprocessed functional data of each participant, which generates p-values for spikes (DVARS) indicating volume-wise severe intensity dropouts. Participants with dropouts in more than 10% of the 300 volumes were excluded (*n* =− 8). Lastly, the “check sample homogeneity was performed using standard deviation across sample” function analysis provided by the CAT12 toolbox (Gaser & Dahnke, 2016) to check for potential misalignments. Participants detected as outlier were manually checked and excluded as the individual mean AROMA functional image did not align to the MNI152 template (*n* =− 8).

### M.3 Structural connectome and functional connectome construction

#### M.3.1 Parcellation

We parceled the whole-brain into 100 discrete and non overlapping regions using the predefined 17-network parcellation scheme by (Schaefer et al., 2018). The components provided by the 17-network parcellation can be allocated to the 7-network scheme, the latter being associated with seven distinct behavioral systems: visual, sensorimotor, limbic, frontoparietal, default-mode, dorsal and ventral-attention network (S. M. Smith et al., 2009; Thomas Yeo et al., 2011).

#### M.3.2. Structural Connectome

For structural connectivity (SC), the parcellation template first had to be warped to individual diffusion space. This was done by combining the non-linear warps of the spatial T1 registration to MNI152 and the distortion correction with the APMs. Since streamlines are generated seeding from the gray-white matter interface and the predefined parcellation scheme only covers cortical gray matter, the template was expanded adding voxels towards the gray-white matter boundary so that all regions also include the seeding points. To increase the biological accuracy of SC, the SIFT-2 method was applied (R. E. Smith et al., 2015). Here, each streamline is weighted with an estimate of its effective cross-sectional area, so that the streamline density matches the white matter fiber density computed directly from the diffusion signal. Based on the estimated streamlines, we calculated the proportion of streamlines between regions (Bassett et al., 2018; Murphy et al., 2020) resulting in a 100×100 adjacency matrix *w_ij_*. The brain was then represented as a whole-brain connectome, in which the parcels are considered as nodes and the structural connections as edges *w_ij_* connecting node i and j. By means of CSD, *w_ij_* was computed as a density of streamlines between two regions by also correcting for the changing cross-sectional area. All 649 SC adjacency matrices (*Regions* × *Regions*) were normalized by the maximum of the entire 1000BD cohort.

#### M.3.3 Functional Connectome

For functional connectivity, we computed both, static FC as well as FC dynamics (FCD). For each of the 100 regions, mean fMRI time-series (BOLD signals) were extracted by averaging the functional activity of all voxels belonging to this region. For static FC, the connectivity (edges) between regions (nodes) was estimated by correlating the mean time-series using the Pearson’s correlation coefficient (static PCC or sPCC) resulting in a 100×100 FC matrix for each participant (Figure 3). Noteworthily, to make our findings comparable to previous work and since negative connectivity values are discussed ambiguously (Chan et al., 2014; Stumme et al., 2020), we only considered the positive edges in the FC and set negative correlation values to zero.

FCD is the dynamic or time-variant representation of FC and reflects the fluctuation of the covariance matrix over time. FCD was quantified by means of a sliding window Pearson correlation coefficient (dynamic PCC or dPCC). Following the procedure in (Battaglia et al., 2020), we estimated the FC for a 40 sec window with maximum overlap (slide step set to 1 sample) to obtain a stream or array of FC matrices that spread over time. The length of the window was set as a trade-off between a lower sampling FC variability and a sufficient sensitivity to detect temporal FC transients (Jia et al., 2017; Lurie et al., 2020). The intrinsic dynamics of the stream was measured as the combinatorial similarity among the matrices of the stream of FCs (Figure 2.E). This match is defined as the Pearson correlation between FC at time or slice *t*_1_ and FC at time or slice *t*_2_ according to the formula

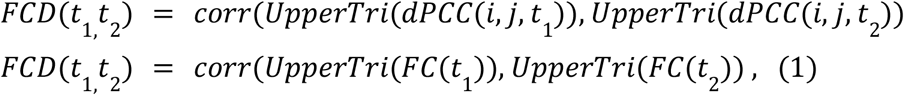

which gives a *timeXtime* matrix known as the FCD Matrix. To assess whether the BOLD time series might have state transitions between high and low activity, we evaluated the switching behavior of BOLD as described in (Hansen et al., 2015; Rabuffo et al., 2021) by employing a compressed metric to assess this transitioning behavior. Therefore, we defined the switching index (SI) or FCD variance as the variance of the upper triangular part of the FCD matrix (Figure 3) or its Frobenius Norm:

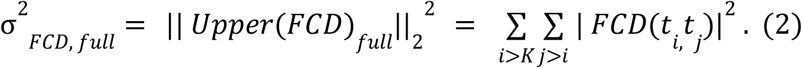

The diagonal order K of the upper triangular matrix was set equal to dPCC window length (converted in samples) to correct for the overlap.

### M.4 The virtual aging brain pipeline

For the investigation of the causal link between white matter degeneration and functional changes and its association with the heterogeneity of cognitive decline, we performed a two-fold analysis. In the first part, we addressed the questions whether realistic brain function can be modeled via a simulated SC decline. After we verified that the presence of a decline in the interhemispheric connectomes in empirical SC dataset, we designed virtual brains, brain network models constrained by the imaging data of SC, to predict brain function from the empirical SC dataset (*SC_emp_*) as well as one virtually aged participant (*SC*_α_) (M.4.1) and again, compared the derived (in-silico) functional data between each other as well as to the empirical (in-vivo) functional data (M.4.2) to ensure the SC changes of the virtual brains could reproduce age-related functional changes. Confident about the model performance, we then, in the second part, inspected the model derived output variable (G-coupling index) characterizing the tethering between SC and FC and related it to age, sex and cognitive performance (M.4.3).

#### M.4.1 Predicting functional connectivity: The Connectome-based Brain Network Model

The approach to virtually age a structural connectome provides the basis to test whether a specific modification in SC affects brain function in a causal sense. Therefore, as a preliminary investigation, we inspected the empirical SC datasets with regards to age and specifically tested whether the previously reported strong age-related interhemispheric SC decline (Jockwitz et al., 2019; Puxeddu et al., 2020; Roland et al., 2017; Zhao et al., 2015) is also applicable to the current dataset of older adults (see Figure 2).

Furthermore, we performed a virtual aging process on the 50 youngest participants (mean age, age-range) by homogeneously decreasing the interhemispheric connections. In particular, we applied a mask on the empirical connectome *W*_0_ as follows:

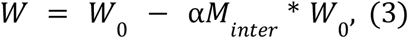

where *M_inter_* is the antidiagonal mask, α is the normalized intensity of decrease and * represents an element-wise product (see Figure 2). The α parameter was spanned in the range [0-1] with sampling interval 0.05 resulting in 20 virtually aged structural connectomes per subject (SC).

The α-masking approach was used to assess the direct impact of the simulated interhemispheric SC decrease on the simulated functional data. To assess the effect of age-related interhemispheric SC degeneration on the brain dynamics in a causal sense, we applied the brain network model in two scenarios. We simulated resting-state activity via virtual brains that couples neural mass (NM) models described in (Montbrió et al., 2015) through the weighted edges in the SC matrix of all participants (n = 649) (Figure 2.A) or via the virtually-aged or α-masked matrices of the 50 youngest subjects (Figure 2.B). The nodal dynamics is a mean-field representation of an ensemble of infinite all-to-all connected quadratic-integrate-and-fire (QIF) neurons at the thermodynamic limit. Following (Montbrió et al., 2015; Rabuffo et al., 2021) and assuming that input currents to the mass system were distributed according to a Lorentzian distribution, the dynamics of each region were given by the following equations:

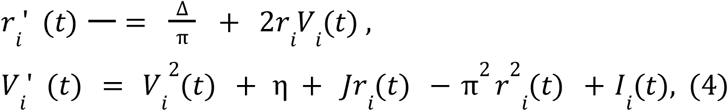

where the variable *r_i_* (*t*) is the population firing rate, while *V_i_*(*t*) is the average membrane potential of the mass. The parameter *J* represents the average synaptic weight, while the parameter η and Δ represent the average excitability of the mass neurons and the heterogeneous noise spread, respectively. These two parameters represent the center and the half width of the Lorentzian distribution of the input currents. The values of these three parameters were set to *J* = 14.5, η = −4.6, Δ = 0.7 in order to obtain a bistable regime, meaning a down-state fixed point and an up-state stable focus in the phase space (Rabuffo et al., 2021). The bistability is a fundamental property to ensure a switching behavior in the data, that has been considered representative of realistic dynamics of real data (Hansen et al., 2015; Rabuffo et al., 2021). The dynamics of a node can arise by the oscillation between these two points (the down-state and the up-state) in the phase space thanks to current *I_i_*(*t*). We could then model a brain network model by *I_i_*(*t*) with:

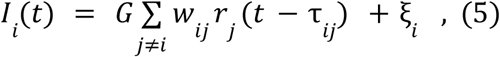

and obtaining the coupled differential equation:

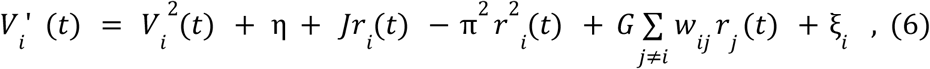

where *r_j_*(*t*) is the firing rate coming from other regions weighted by the structural connectivity edges *w_ij_*. The variable ξ_*i*_ represents the noise that models stochastic external factors and allows the region i to oscillate between the two states. The parameter *G* is the global parameter that modulates the overall impact of SC on the state dynamics of each region and the ξ_*i*_ is the noise variable that ensures the oscillations between the up-state and down-state and is distributed according to a Gaussian distribution with a variance set to σ.

For each connectome, the brain network model was fitted by optimizing parameters *G* modulation index and noise variance σ^2^ to maximize the SI or σ^2^_*FCD, full*_ of the simulated BOLD data (see Figure 3). SI is an indicator for the metastability or switching behavior of functional brain networks and can be used as a measurement for the realistic evaluation of brain function. It has been analyzed in various studies related to both, whole-brain modeling (Deco et al., n.d.; Hansen et al., 2015; Rabuffo et al., 2021) and aging (Battaglia et al., 2020). In fact, previous results indicate simulated data which was created by the enhancement of a metastable or switching behavior and tuning model parameters have been considered to provide more realistic dynamics than those minimizing the distance between the empirical FC and the simulated one (Courtiol et al., 2020; Deco et al., n.d.; Hansen et al., 2015). The capacity to replicate empirical data does not only entail the ability to reproduce a specific property of the functional data, but reproduce a wider range of features or summary statistics, such as static FC and other FCD properties. Specifically, for each dataset we conducted a parameter sweep or grid search for the global parameter in the range *G* = [1. 5 − 3.2] with sampling interval Δ*G* = 0.05 and the noise variance in the range σ^2^ = [0.01 − 0.05] with Δ σ^2^ = 0.01 to determine the optimal couple (*G*, σ^2^) for which SI is maximal. The direct numerical integration for the nodal equations (2) was performed via Heun-stochastic integration implemented in the Virtual brain open-source code. For each region, we integrated the equations with variable time steps (from 0.005 ms to 0.0005 ms) for 5 minutes and we downsampled the variables *r_i_* (*t*), *V_i_*(*t*) to the sampling frequency *fs* = 100 *Hz*. The time step was variable to adapt the integration process and avoid numerical errors or instabilities and the neural mass field was filtered via the Balloon-Windkessel model (Friston et al., 2000) to emulate fMRI time series with repetition time TR set to 2000 ms.

For each model fit (the 649 empirical and 20*50=1000 virtually aged connectomes), we obtained a *G* modulation index which is representative of how much the SC influences the brain dynamics. Before investigating if the SC-FC causal link was related to behavioral factors such as age, sex and cognitive performance, we compared the in-silico brain dynamics to in-vivo data assuring that a variety of dynamic and static properties of functional covariates are indeed realistic.

#### M.4.2 Comparing estimated in-silico and in-vivo functional data

To ensure that the maximization of the SI indeed provides realistic functional dynamics, we estimated and compared a set of dynamic and static FC properties in the empirical data, in the age-simulated data and in the alpha-simulated data (or virtually aged data). In particular, we focused on functional biomarkers that have previously been shown to be biologically related to white-matter degeneration at microstructure level and to the cross-hemispheric SC topology (Mollink et al., 2019; Roland et al., 2017).

Specifically, we computed the average homotopic FC strength, the FCD variance difference, the standard deviation of the interhemispheric FC stream. The average homotopic FC strength is defined as the average sum of the specular interhemispheric connections, as follows:

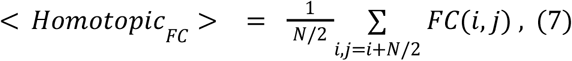

where the averaged *FC*(*i, j*) entries represent the element of order 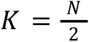. This definition is derived by the microstructure-function investigation by (Mollink et al., 2019). We derived both the average of all homotopic edges or the average of edges connecting homotopic regions within a certain rsFC network. To extend the SI towards an interhemispheric topology, we computed the FCD variance difference as the difference between the SI of interhemispheric FCD matrix and the SI of the full FCD matrix, as follows:

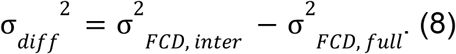

The σ^2^_*FCD, inter*_ was computed by considering only the interhemispheric edges of dPCC in each window and therefore in the similarity computation (see Figure 3). The variance is then obtained as the Frobenius norm of the upper triangular part of the interhemispheric FCD matrix:

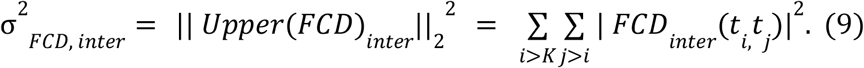

Similarly, one can compute the standard deviation of the interhemispheric FC stream as the variability of the interhemispheric FC edges of the FC matrix stream obtained with the windowing scheme. By unraveling the FC matrix in each slice, we obtained a vector ‘FC edges’ for each time point or *edgesXtime* matrix. We derived the standard deviation of all interhemispheric edges’ time-courses as a representative index of the connectivity oscillations across hemispheres or the stability of the metaconnectivity (Esfahlani et al., 2021; Faskowitz et al., 2020).

To test whether we could actually reproduce the dynamical and statical properties of FC, we estimated linear trends of the homotopic FC and FCD variance difference features in the functional empirical data, the age-simulated data and the alpha-simulated data. Specifically, we performed Pearson correlations between the functional summary statistics and age or the alpha masking variable, respectively. To evaluate whether the masking had a nonlinear effect on the functional features trends, we computed both, a linear fit and quadratic fit for the virtually aged data.

#### M.4.3 SC-FC tethering and its association to age, cognitive performance and sex

The model-derived tethering between SC and FC (*G* modulation index) was then statistically related to both demographic and behavioral factors.

First, we performed an ANCOVA indicating whether the *G* index is overall age- or sex-related (corrected for education). Furthermore, to analyze age-related trajectories of the SC-FC linkage, we performed partial correlations between the *G* modulation index and age (corrected for sex and education). This was repeated after stratifying the group by sex, and the partial correlations (corrected for education) were statistically compared (Fishers-Z) indicating whether age-related *G* modulation index changes are different in males as compared to females. Lastly, we performed a Pearson’s correlation to test the relation between the *G* modulation index of the virtually aged dataset and the alpha-variable.

According to various aging theories (Festini et al., 2018; Reuter-Lorenz & Park, 2014), changes in the SC-FC tethering might explain the variability of cognitive performance during the aging process. To test this hypothesis, we performed ordinary-least squares (OLS) multivariate regressions for each of the 16 cognitive scores, testing the relation between the cognitive performance and the G-coupling index, while correcting for age, sex and education. We specified the model as:

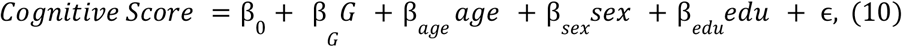

Here, results were considered significant at *p* ≤ 0.05, Bonferroni corrected for the number of performed models (*n* = 16 cognitive scores, *p_cor_* = *p*_β_ * 16).

Moreover, for cognitive performance scores showing a significant relation to the *G* modulation index, we were interested whether the strength of performance determines a different age-related trajectory in the *G* index. Specifically, we split the whole group by the performance median into a low performing (LP) and a high performing group (HP) and tested whether the *G* modulation index is significantly different between groups (ANCOVA corrected for sex, education and age). We also calculated partial correlations between age and the *G* modulation index (corrected for sex and education) and compared these between groups (Fishers-Z). Post-hoc, based on a depicted non-linear trend of the correlation between age and the *G* modulation index in particularly the low performing subjects, we split the groups by the median age (67 years) into a younger and older group. Partial correlations between age and the *G* modulation index (corrected for sex and education) were repeated group-wise (for low and high performers in young and older subjects) and subsequently compared between low and high performing groups using Fishers-Z (see Figure 4).

### M.5 Simulation-based inference

The global modulation parameter *G* of the brain network model was also estimated by simulated-based inference (SBI) tailored to Bayes rule in order to retrieve the model parameter space compatible with the empirical data (Cranmer et al., 2020; Gonçalves et al., 2020). Statistical inference consists of the automatic identification of possible parameters θ via the likelihood function *p*(*X*|θ), which quantifies the probability that a given set θ generates the vector of raw data or low-dimensional data features *X*. The Bayesian estimation consists in *p*(*X*|θ) = *p*(*X*|θ) *p*(θ), where *p*(*X*|θ) is the posterior that quantifies the probabilistic consistency between the selected parameter space and fitted empirical data. In order to determine the likelihood function for a high-dimensional mechanistic model such as the brain network model in this study, we used the sequential neural posterior estimation (SNPE) to approximate the distribution *p*(*X*|θ) with a flexible neural density estimation trained on low-dimensional data features (Gonçalves et al., 2020). SNPE is based on the training of a deep neural network, which allows to directly approximate all posteriors from ad-hoc data features, where the calculation of likelihood function is analytically or computationally intractable. Based on a “budget” of simulations obtained by a mechanistic model and a specific set of (low-dimensional) summary statistics obtained from the generated data, the weights of the network are optimized by loss minimization via masked autoregressive flow (MAF) bypassing the need for the highly computational intensive Markov-Chain Monte Carlo sampling (Papamakarios et al., 2017). Therefore, this approach is considered as a simulation-based inference approach that is amortized: after an upfront computational cost at the simulation and training steps, a new data set of the subject can be fitted efficiently by a single forward pass of the empirical data through the neural network without the computational overhead for further simulations at the inference step (Cranmer et al., 2020; Gonçalves et al., 2020). In this study, a “budget” of 2000 simulations was obtained by the model in equation (2) and the network was trained on a matrix *X*, which contained the following three features or summary data for each simulation: average homotopic FC strength, the difference between the whole-brain FCD and the interhemispheric FCD, the standard deviation of the interhemispheric FC stream. The simulation were performed with a *G* distributed according to a uniform prior (truncated between 1.5 and 3.2), such that the final posterior estimation resulted in: *p*(*G_SBI_*|*X_emprical_*) = *p*(*X_stimulated_*|*G_SBI_*). As highlighted by the formula, the SNPE was repeated on each subject, trained on the simulated data and the final estimation of *G* was obtained by maximizing the log likelihood of the features of the empirical BOLD data under the model. The SNPE provides the statistical relation between G and the empirical data and the final estimate of GSBI was considered as the mean value of the estimated posterior. For each individual SBI, the noise variance was set to the optimized value obtained via the parameter sweep. The quality of SNPE was evaluated as the Pearson correlation between *G_SBI_* and *G_Sweep_* as well as the trend of *G_SBI_* with age. We also checked the uncertainty of the estimations encoded in posterior distribution for each subject. The SNPE estimation was also validated using the in-silico data generated by TVB for one subject, as the estimated posterior distributions encompass the ground truth values used in the simulation (see Figure S2 in the supplements).

## Code availability

The code of the VAB pipeline will be available at the following link: https://github.com/ins-amu/virtual_aging_brain

## Supplementary materials

**Supplementary Table 1:**
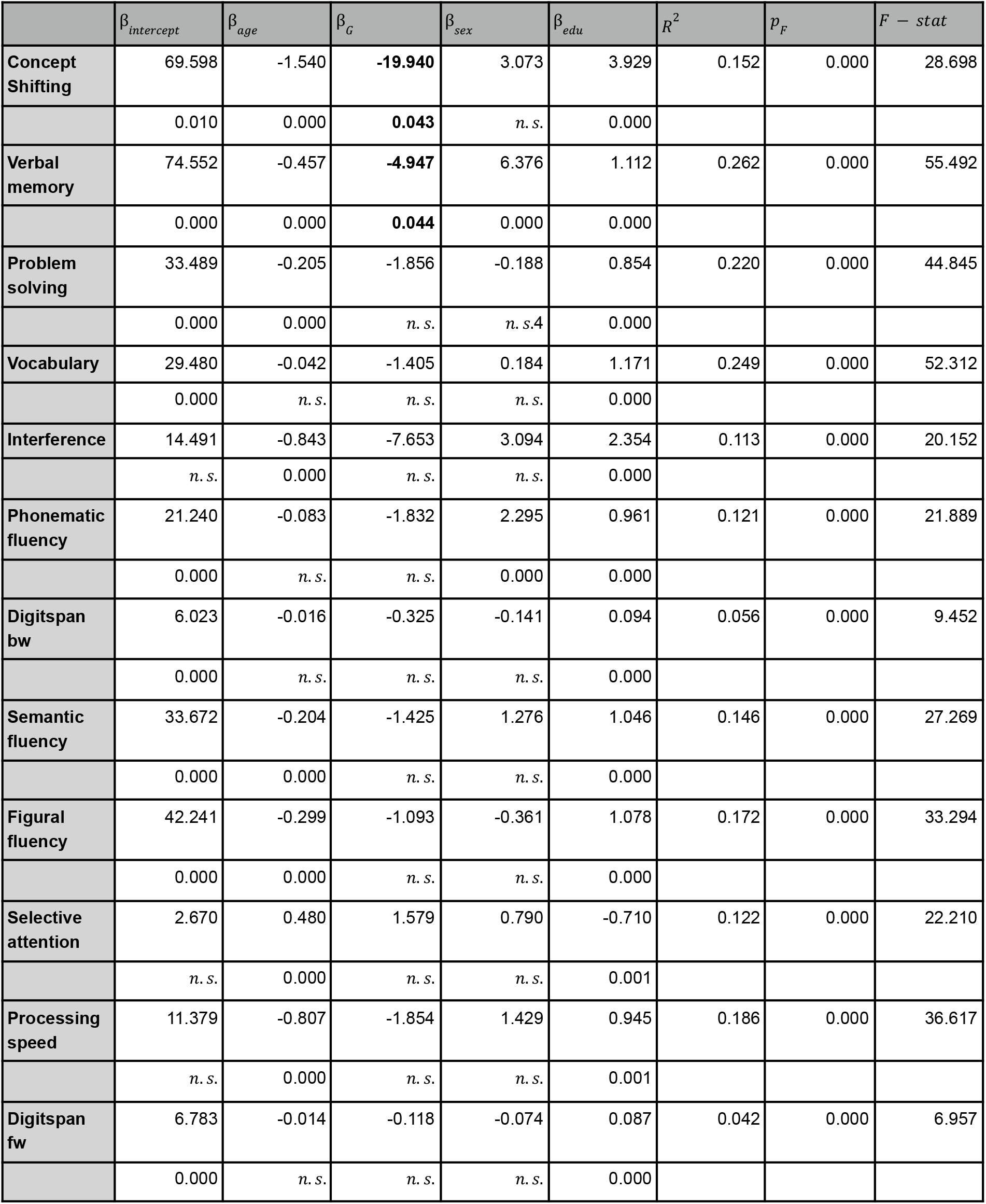
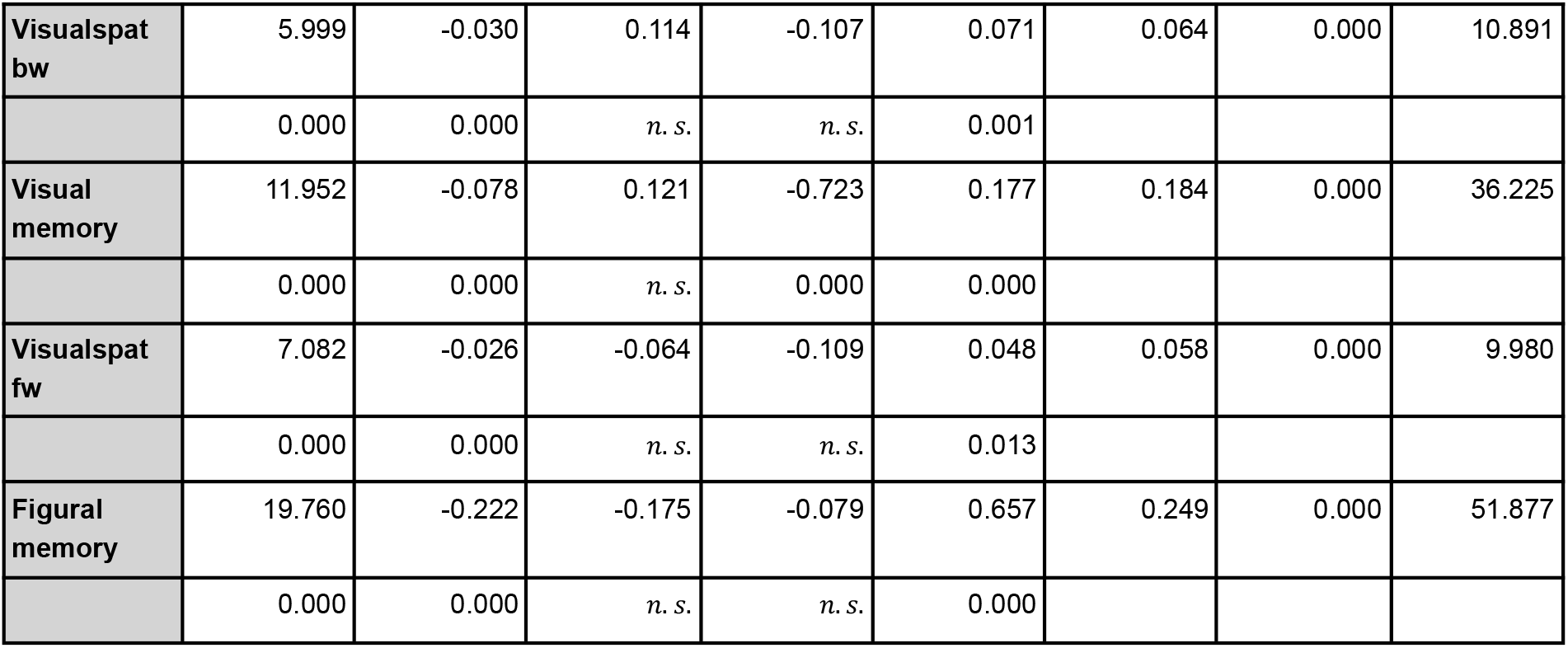
Results of the ordinary least squares regression to link cognitive scores as dependent variables with age, sex and education as indipendent ones. The first five columns represent the coefficients of the model and the associated p-value, while the latter three are the coefficient of determination *R*^2^, the p-value of the F-test (*p_F_*) and the F-statistics (*F* − *stat*), respectively.

| | $\beta_{intercept}$ | $\beta_{age}$ | $\beta_G$ | $\beta_{sex}$ | $\beta_{edu}$ | $R^2$ | $p_F$ | $F - stat$ |
| --- | --- | --- | --- | --- | --- | --- | --- | --- |
| **Concept Shifting** | 69.598 | -1.540 | **-19.940** | 3.073 | 3.929 | 0.152 | 0.000 | 28.698 |
|  | 0.010 | 0.000 | **0.043** | *n. s.* | 0.000 |  |  |  |
| **Verbal memory** | 74.552 | -0.457 | **-4.947** | 6.376 | 1.112 | 0.262 | 0.000 | 55.492 |
|  | 0.000 | 0.000 | **0.044** | 0.000 | 0.000 |  |  |  |
| **Problem solving** | 33.489 | -0.205 | -1.856 | -0.188 | 0.854 | 0.220 | 0.000 | 44.845 |
|  | 0.000 | 0.000 | *n. s.* | *n. s.* | 0.000 |  |  |  |
| **Vocabulary** | 29.480 | -0.042 | -1.405 | 0.184 | 1.171 | 0.249 | 0.000 | 52.312 |
|  | 0.000 | *n. s.* | *n. s.* | *n. s.* | 0.000 |  |  |  |
| **Interference** | 14.491 | -0.843 | -7.653 | 3.094 | 2.354 | 0.113 | 0.000 | 20.152 |
|  | *n. s.* | 0.000 | *n. s.* | *n. s.* | 0.000 |  |  |  |
| **Phonematic fluency** | 21.240 | -0.083 | -1.832 | 2.295 | 0.961 | 0.121 | 0.000 | 21.889 |
|  | 0.000 | *n. s.* | *n. s.* | 0.000 | 0.000 |  |  |  |
| **Digitspan bw** | 6.023 | -0.016 | -0.325 | -0.141 | 0.094 | 0.056 | 0.000 | 9.452 |
|  | 0.000 | *n. s.* | *n. s.* | *n. s.* | 0.000 |  |  |  |
| **Semantic fluency** | 33.672 | -0.204 | -1.425 | 1.276 | 1.046 | 0.146 | 0.000 | 27.269 |
|  | 0.000 | 0.000 | *n. s.* | *n. s.* | 0.000 |  |  |  |
| **Figural fluency** | 42.241 | -0.299 | -1.093 | -0.361 | 1.078 | 0.172 | 0.000 | 33.294 |
|  | 0.000 | 0.000 | *n. s.* | *n. s.* | 0.000 |  |  |  |
| **Selective attention** | 2.670 | 0.480 | 1.579 | 0.790 | -0.710 | 0.122 | 0.000 | 22.210 |
|  | *n. s.* | 0.000 | *n. s.* | *n. s.* | 0.001 |  |  |  |
| **Processing speed** | 11.379 | -0.807 | -1.854 | 1.429 | 0.945 | 0.186 | 0.000 | 36.617 |
|  | *n. s.* | 0.000 | *n. s.* | *n. s.* | 0.001 |  |  |  |
| **Digitspan fw** | 6.783 | -0.014 | -0.118 | -0.074 | 0.087 | 0.042 | 0.000 | 6.957 |
|  | 0.000 | *n. s.* | *n. s.* | *n. s.* | 0.000 |  |  |  |
| **Visualspat bw** | 5.999 | -0.030 | 0.114 | -0.107 | 0.071 | 0.064 | 0.000 | 10.891 |
|  | 0.000 | 0.000 | *n. s.* | *n. s.* | 0.001 |  |  |  |
| **Visual memory** | 11.952 | -0.078 | 0.121 | -0.723 | 0.177 | 0.184 | 0.000 | 36.225 |
|  | 0.000 | 0.000 | *n. s.* | 0.000 | 0.000 |  |  |  |
| **Visualspat fw** | 7.082 | -0.026 | -0.064 | -0.109 | 0.048 | 0.058 | 0.000 | 9.980 |
|  | 0.000 | 0.000 | *n. s.* | *n. s.* | 0.013 |  |  |  |
| **Figural memory** | 19.760 | -0.222 | -0.175 | -0.079 | 0.657 | 0.249 | 0.000 | 51.877 |
|  | 0.000 | 0.000 | *n. s.* | *n. s.* | 0.000 |  |  |  |

**Supplementary Table 2:** Description and Median (25% percentile - 75% percentile) values of the 1000BRAINS cognitive scores (Caspers et al., 2014).

|  | Median | q25 | q75 |
| --- | --- | --- | --- |
| **Visualspat fw** | 5 | 5 | 6 |
| **Visualspat bw** | 5 | 4 | 5 |
| **Visual memory** | 8 | 6 | 9 |
| **Figural fluency** | 26 | 21 | 31 |
| **Problem solving** | 21 | 17 | 24 |
| **interference** | -38.42 | -49.885 | -30.055 |
| **Concept shifting** | -45.235 | -66.79 | -30.0975 |
| **Digitspan bw** | 4 | 4 | 5 |
| **Digitspan fw** | 6 | 5 | 7 |
| **vocabulary** | 31 | 28 | 34 |
| **phonematic_fluency** | 18 | 14 | 23 |
| **semantic_fluency** | 24 | 19 | 29 |
| **Verbal memory** | 43 | 35 | 50 |
| **Figural memory** | 9 | 6 | 12 |
| **Processing speed** | -37.44 | -47.355 | -30.155 |
| **Selective attention** | 32.18 | 27.62 | 38.6525 |

**Figure S1:**
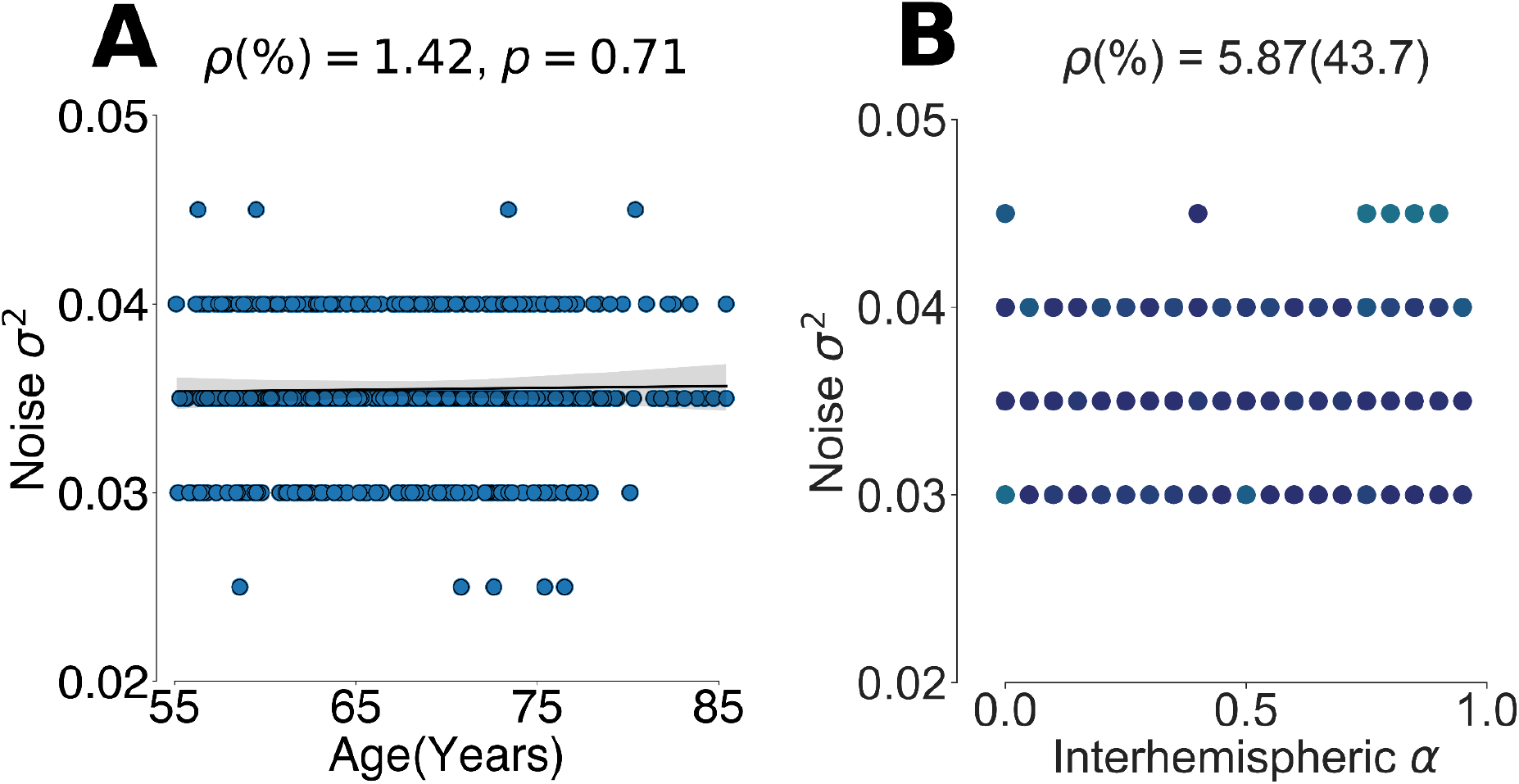
The free parameter Noise variance trend for the empirical connectome based model (*SC_emp_*) in function of age and for the virtually aging process (*SC*_α_) in function of alpha, respectively.

**Figure S2:**
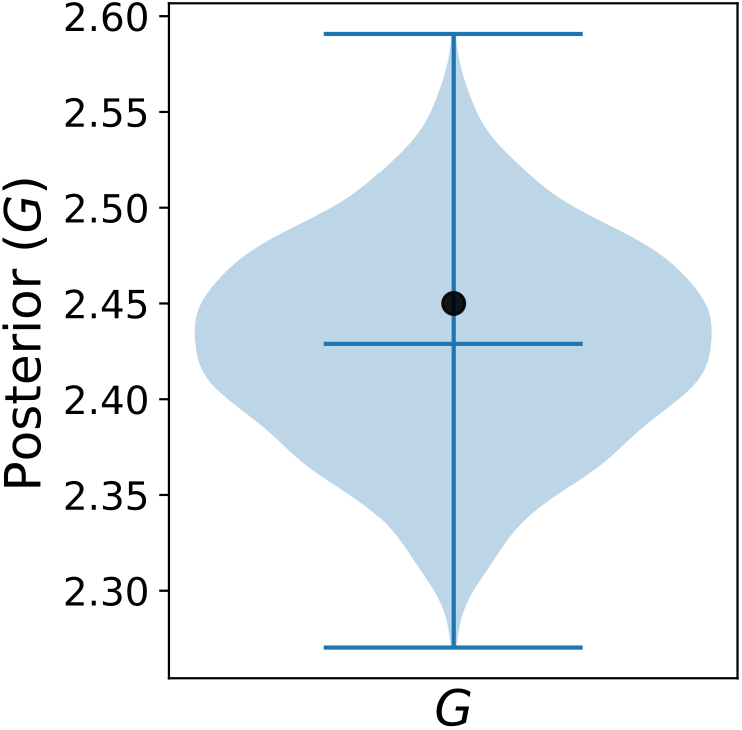
Ground truth testing for SBI approach. The figure shows the posterior estimation of the global modulation G for simulation data whose SC-FC index is known (i.e. ground truth G = 2.45).

**Figure S3:**
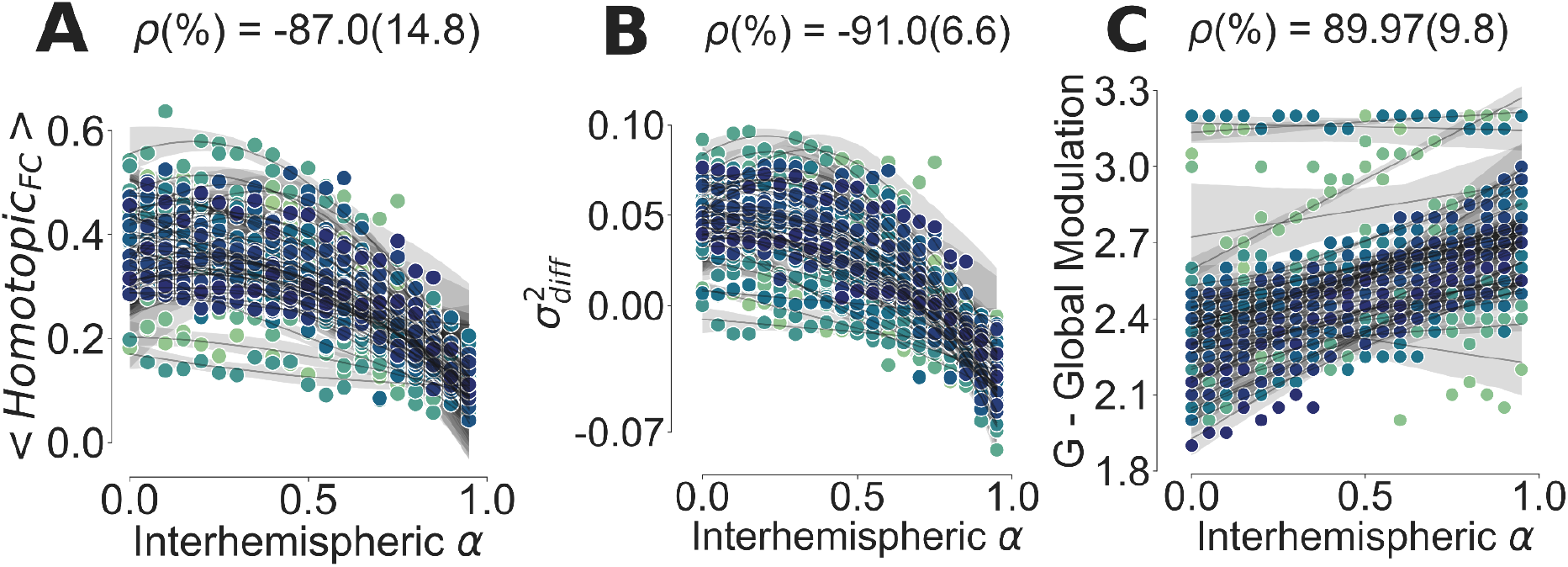
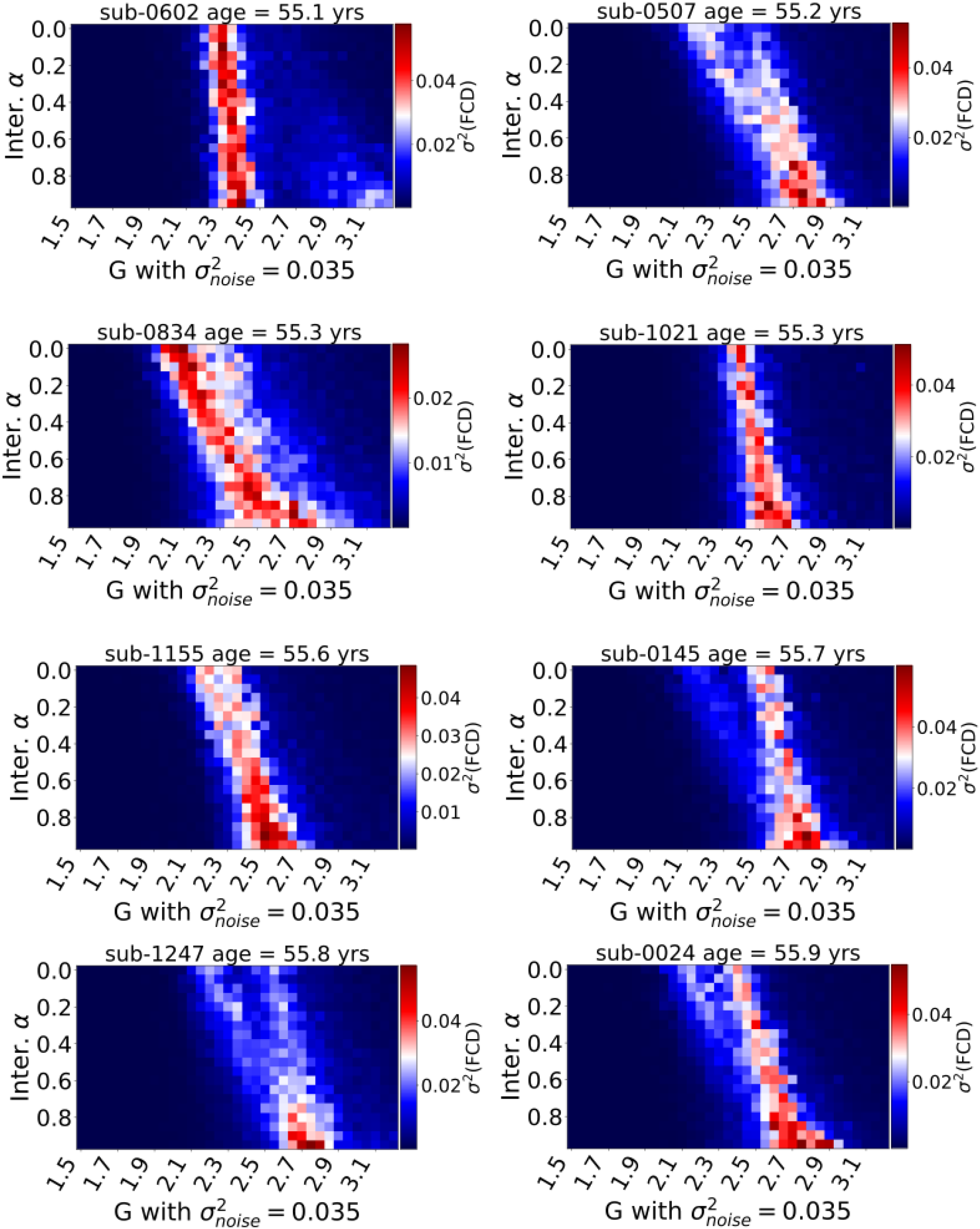
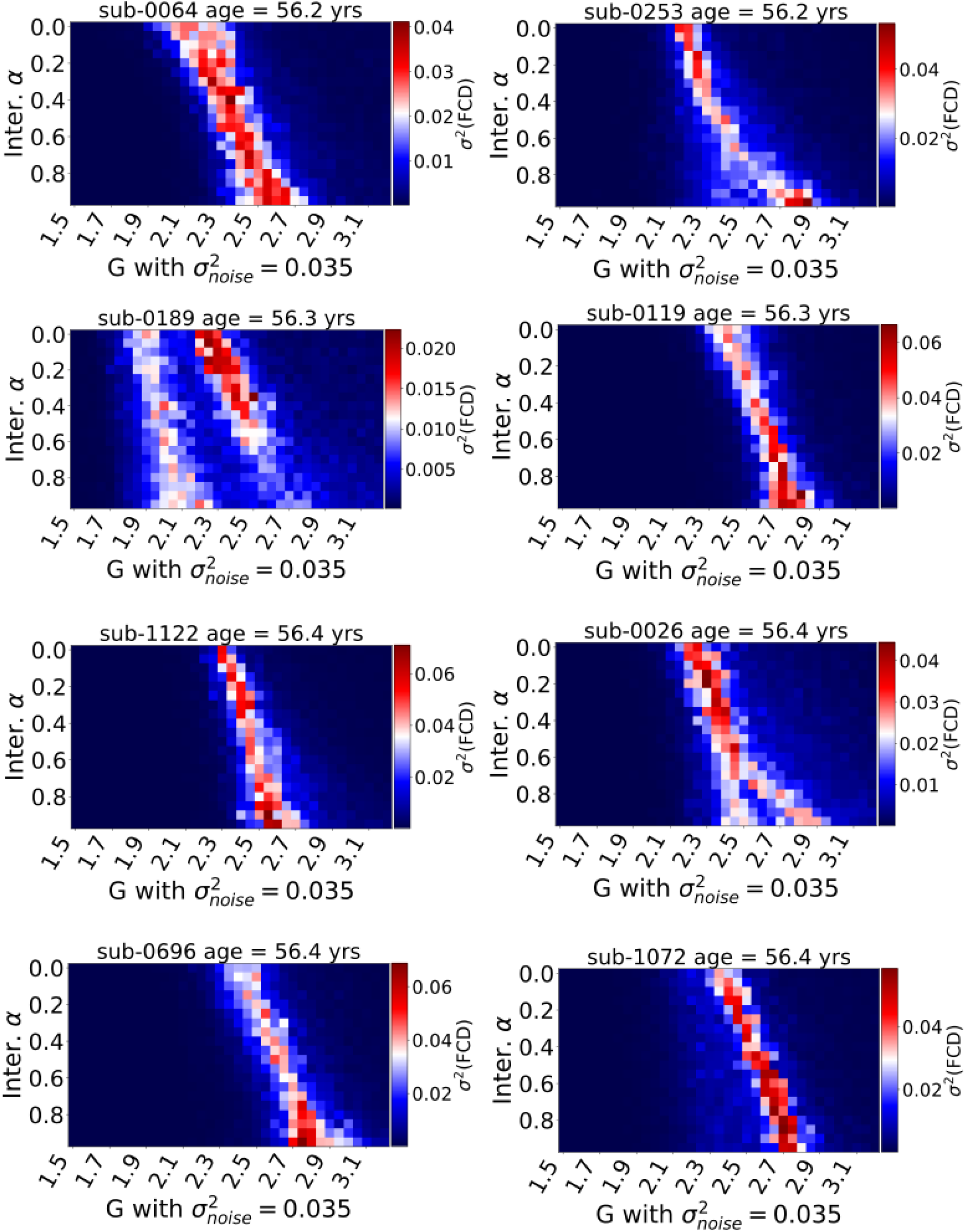
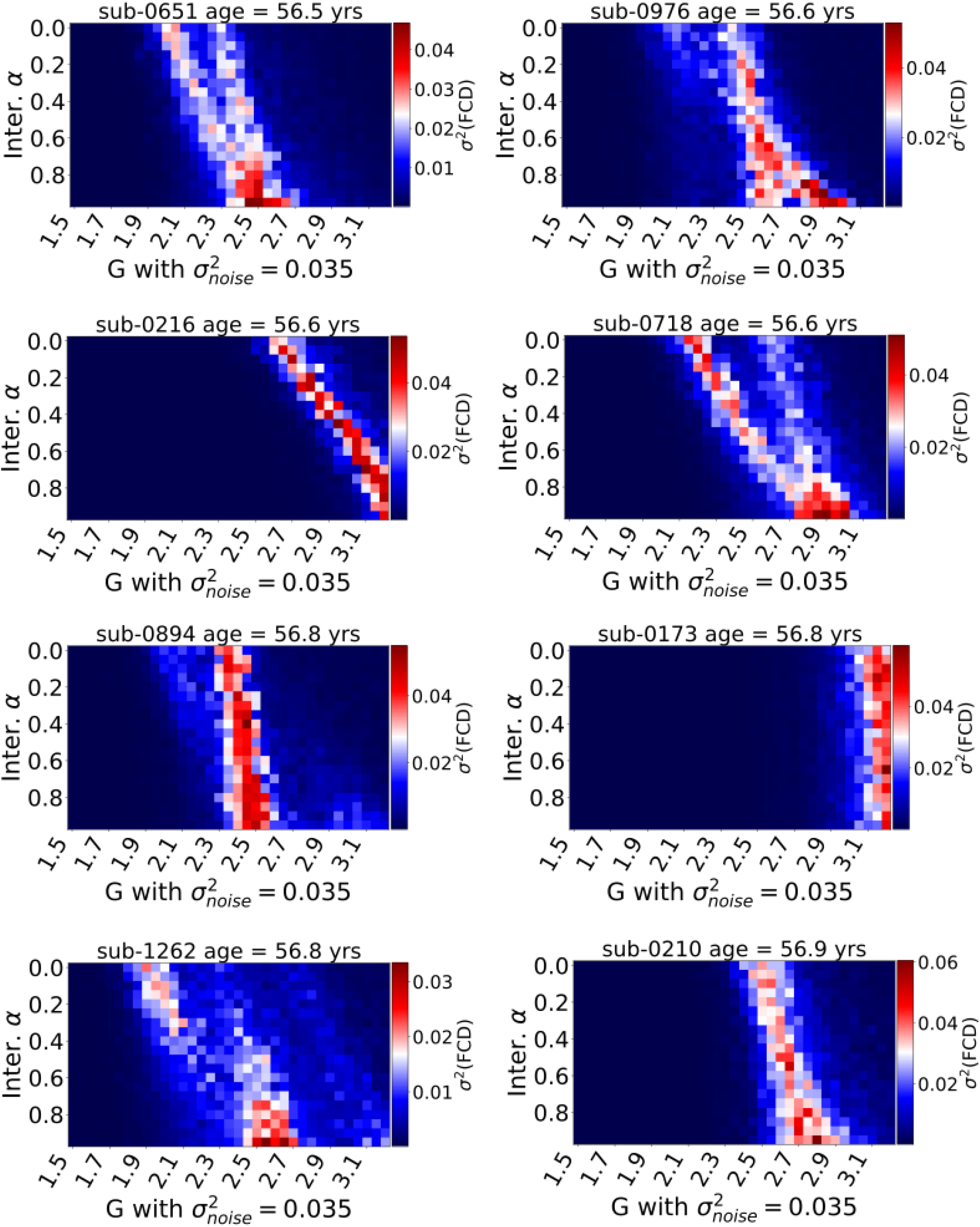
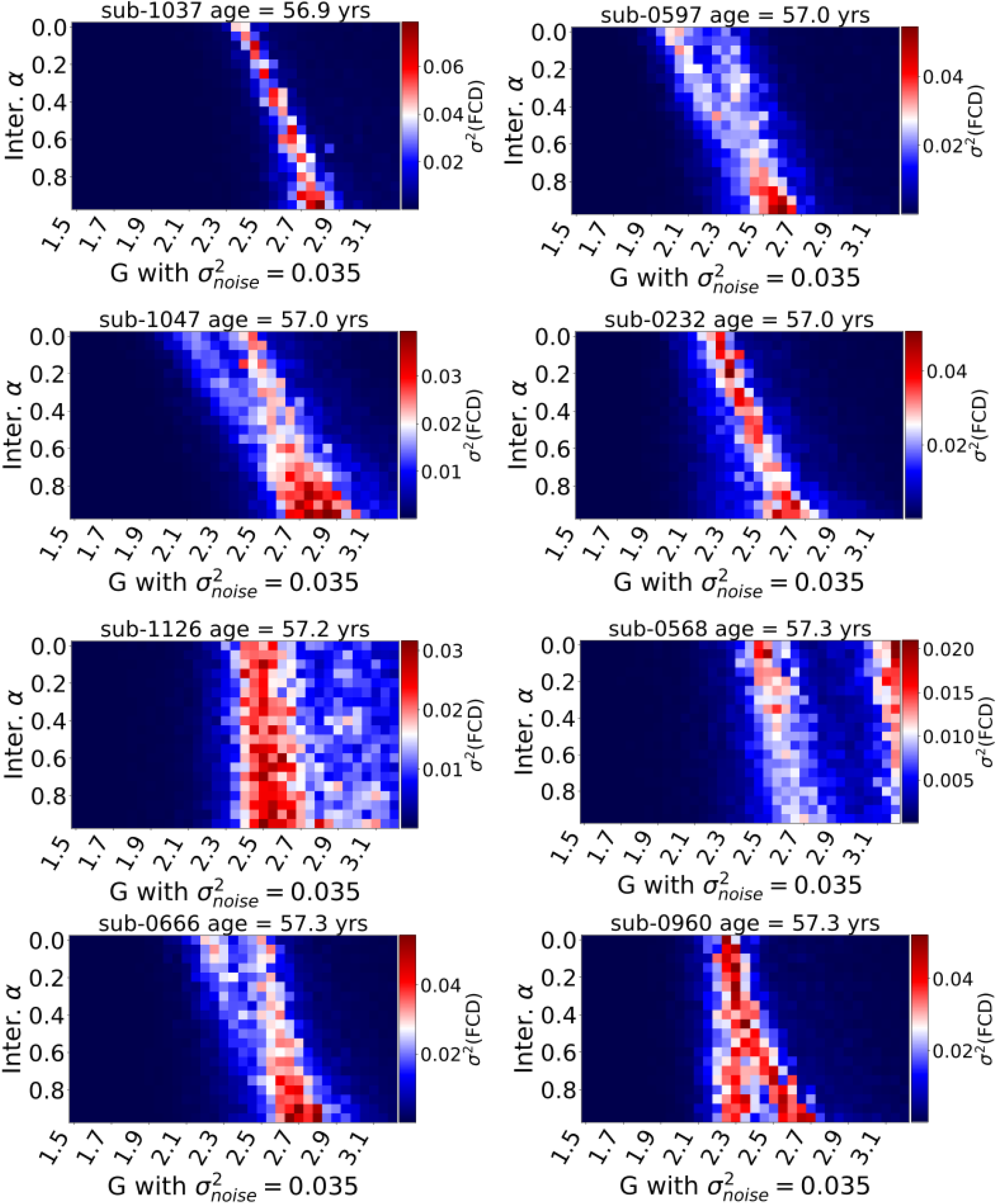
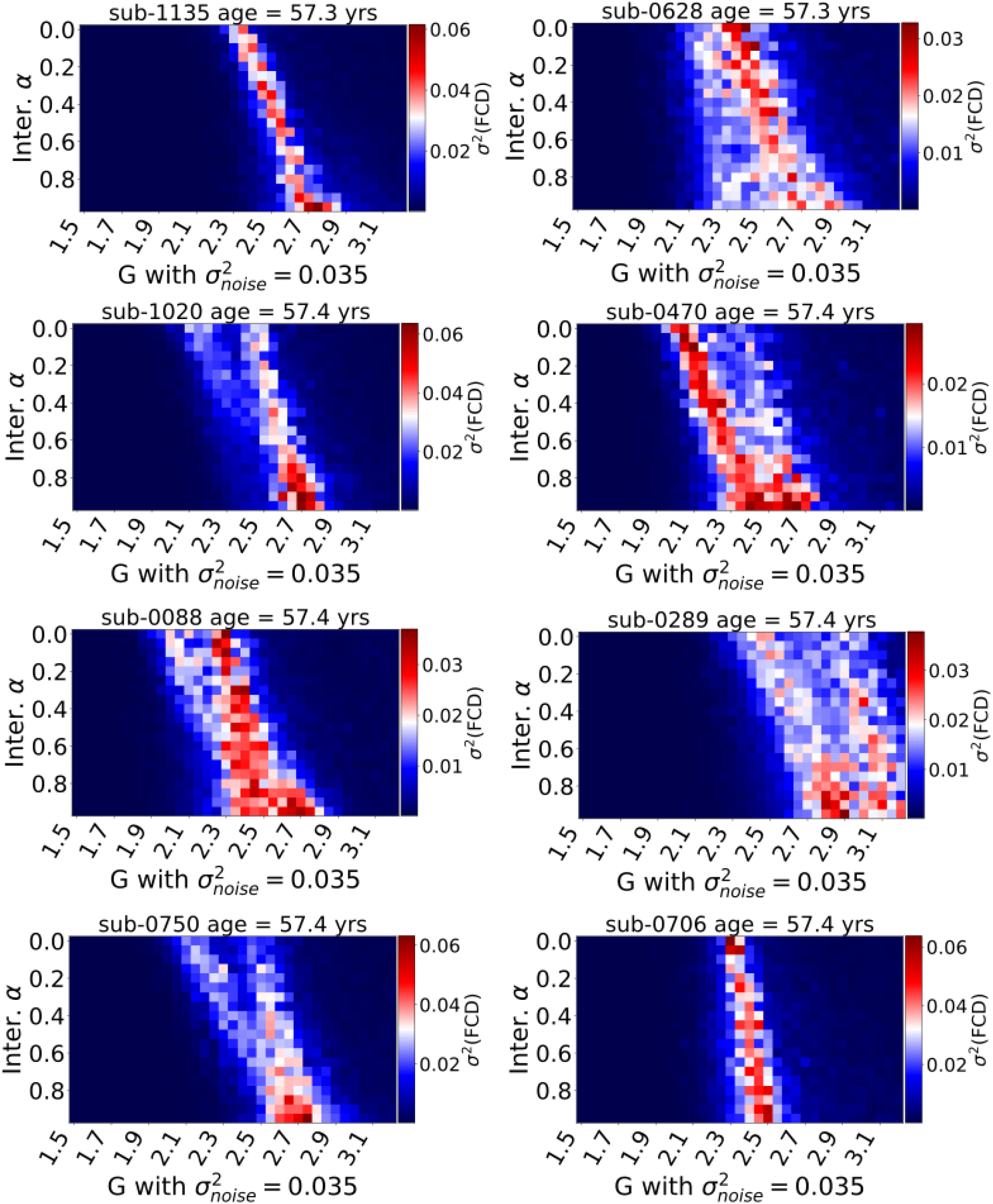
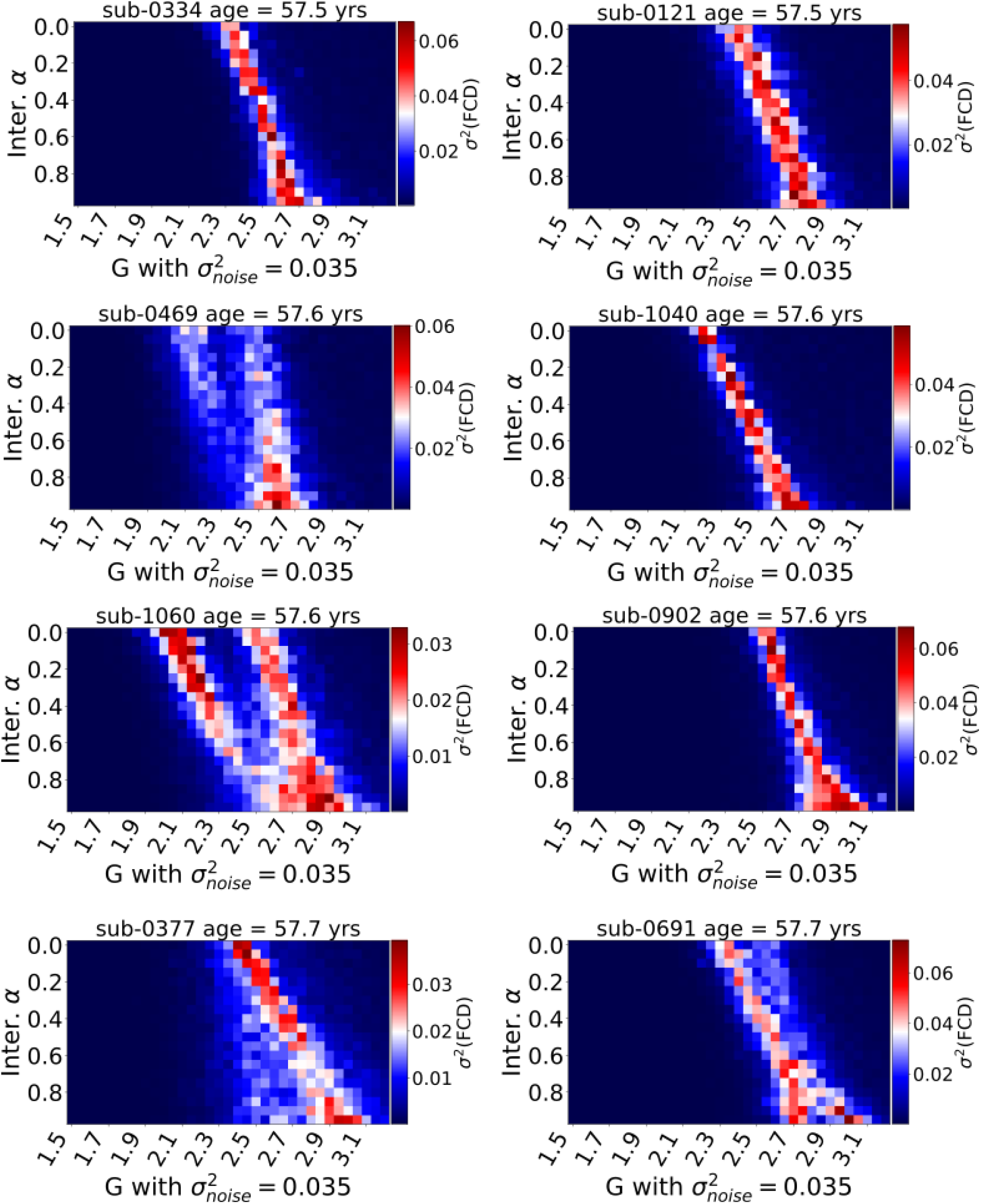
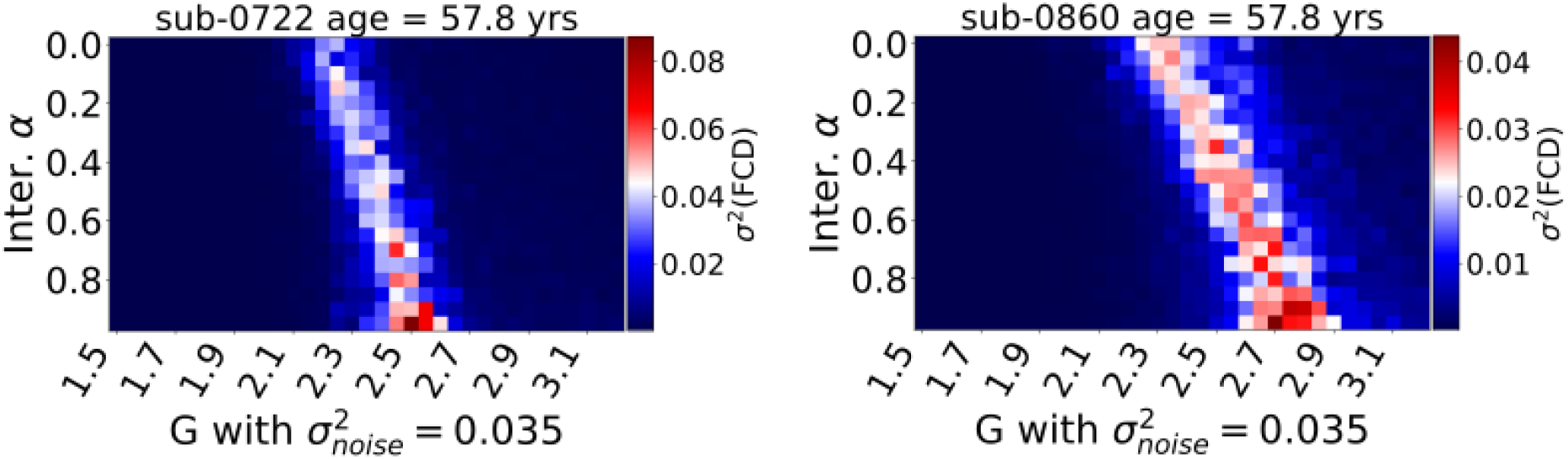
The first two panels show the functional changes of the homotopic FC and FCD variance difference under various degrees of interhemispheric SC degeneration in *SC*_α_ dataset. We also highlighted an individual fit for each of 50 youngest subjects. The latter shows the respective G-modulation index in function of alpha. The following images show the individual SI index (α, *G*) heatmaps for each of the 50 subjects.

